# Crop-photoperiodism model 2.0 for the panicle-initiation date of sorghum and rice that includes daily changes in sunrise and sunset times

**DOI:** 10.1101/2020.11.29.402628

**Authors:** B. Clerget, M. Sidibe, C.S. Bueno, C. Grenier, T. Kawakata, A.J. Domingo, H.L. Layaoen, N.G. Palacios, J.H. Bernal, G. Trouche, J. Chantereau

## Abstract

**Background and Aims:** Daylength determines flowering dates. However, questions remain regarding flowering dates in the natural environment, such as the synchronous flowering of plants sown simultaneously at highly contrasting latitudes. The daily change in sunrise and sunset times is the cue for the flowering of trees and for the synchronisation moulting in birds at the equator. Sunrise and sunset also synchronise the cell circadian clock, which is involved in the regulation of flowering. The goal of this study was to update the photoperiodism model with knowledge acquired since its conception.

**Methods:** A large dataset was gathered, including four 2-year series of monthly sowings of 28 sorghum varieties in Mali and two 1-year series of monthly sowings of eight rice varieties in the Philippines to compare with previously published monthly sowings in Japan and Malaysia, and data from sorghum breeders in France, Nicaragua, and Colombia. An additive linear model of the duration in days to panicle initiation (PI) using day length and daily changes in sunrise and sunset times was implemented.

**Key Results:** Simultaneous with the phyllochron, the duration to PI of field crops acclimated to the mean temperature at seedling emergence within the usual range of mean cropping temperatures. A unique additive linear model combining daylength and daily changes in sunrise and sunset hours was accurately fitted for any type of response in the duration to PI to the sowing date without any temperature input. Once calibrated on a complete and an incomplete monthly sowing series at two tropical latitudes, the model accurately predicted the duration to PI of the concerned varieties from the equatorial to the temperate zone.

**Conclusions:** The results of the updated photoperiodism model strongly supported the hypothesis that photoperiodism could be a combined response to day length and daily changes in sunrise and sunset times.

## INTRODUCTION

In plants and animals, reproductive cycles are synchronised through different means, among which photoperiodism, i.e. a response to the seasonal changes in daylength, plays an essential role (Thomas and Vince-Prue, 1997; Gwinner and Scheuerlein, 1998). This cue, caused by the movement of the earth around the sun, is highly stable across years. After the influence of daylength on the flowering time was understood (Tournois, 1912; Klebs, 1913), many experiments were conducted under either artificial stable daylength or natural changing daylength to characterise the specific response of each species and cultivar. This response to photoperiod is hereafter referred to as photoperiodism. Sorghum and rice have been classified as short days plants in which flowering is proportionally accelerated by short days (Thomas and Vince-Prue, 1997). Vergara and Chang (1985) compiled a large number of such experiments on rice and measured the duration to flowering in a large panel of rice varieties at four stable photoperiods (10, 12, 14, and 16 h light). The results showed substantial variability in the reaction to daylength, from insensitive to highly sensitive. Similar variability was documented in sorghum in 22,500 accessions of the International Crops Research Institute for the Semi-arid Tropics (ICRISAT) genebank sown in fields in Hyderabad, India (17°N) in June and October (Grenier *et al.,* 2001). In tropical areas, photoperiod-sensitivity was also evaluated through the comparison of the durations to flowering in response to monthly sowings conducted over the year (Dore, 1959; Bezot, 1963; Miller *et al.,* 1968; Dingkuhn *et al.,* 1995, 2015; Sié *et al.,* 1998; Clerget *et al.,* 2004). Less commonly, sowings were conducted in a greenhouse over the year at a temperate latitude (Kawakata and Yajima, 1995). In both sorghum and rice species, an annual series of monthly sowings exhibited large ranges in photoperiod sensitivity. Moderately photoperiodic varieties showed quantitative responses to the sowing date, typically modelled through a linear response to daylength (Major, 1980). Highly photoperiodic varieties showed a steep break in the response of the duration to flowering between two sowing dates occurring from January to March, depending on the variety (Miller *et al.,* 1968; Kawakata and Yajima, 1995; Clerget *et al.,* 2004). The response to daylength of this second group of varieties was classified as qualitative photoperiod-sensitive and modelled accordingly using a critical photoperiod threshold, above which flowering was strictly inhibited (Carberry *et al.,* 1992, 2001). In sorghum, it was additionally hypothesised that this threshold decreased as the plant aged because panicle initiation (PI) occurred from July at a higher threshold than the critical threshold that caused the initial flowering inhibition, and that PI immediately occurred when the daylength reached this moving threshold (Folliard *et al.,* 2004; Dingkuhn *et al.,* 2008). Models of photoperiodism generally assume that the crop progresses towards PI by accumulating daily progress equal to the inverse of the predicted duration to PI under the temperature and photoperiod of the day until reaching the value of 1 (Summerfield *et al.,* 1997). This concept of quantitative accumulation was strongly supported by the reciprocal transfer experiments at stable daylength (Collinson *et al.,* 1992; Ellis *et al.,* 1992). Plants were transferred from long to short photoperiod and reciprocally and the duration to PI was a linear combination of the effect of each photoperiod during the time spent in each environment.

A possible effect of decreasing daylength on the triggering of the PI was tested on the dates of PI recorded in monthly plantings of three tropical varieties in Samanko, Mali (12°34′N), but could be successfully parametrised into only two varieties (Clerget *et al.,* 2004). In rice, modern varieties exhibit little photoperiod-sensitivity; thus, no model was developed for the strongly photoperiodic flowering response. Thus, Dingkuhn *et al.* (2015) used the 2008 sorghum Impatience model for highly photoperiod-sensitive rice varieties. All current models are only valid at the latitude where they have been calibrated because photoperiods are different at other latitudes, although Abdulai *et al.* (2012) made a preliminary attempt to adjust the sorghum Impatience model with a latitude effect based on experiments from 11°13′N to 13°15′N.

The effect of the change in photoperiod with latitude on the flowering date has remained an unsolved question. Curtis (1968) showed that six Nigerian varieties sown at three latitudes from 7°30′N to 11°N headed simultaneously despite the difference in the perceived daylengths. In a fourth latitude, 12°N, sowings were delayed by the late onset of the rainy season and discarded. Conversely, Abdulai *et al.* (2012) showed that PI of seven sorghum varieties simultaneously sown in three locations was significantly earlier at 11°13′ than at 12°39′N or 13°25′N. Lafarge (1998) questioned the similar duration to PI of the moderately sensitive sorghum variety E-35-1 simultaneous sown in May in Bamako, Mali (12°39′N) and Montpellier, France (43°39′N), despite the difference of 2 to 3 h in the photoperiods at the two locations.

The cue perceived by plants and animals at equatorial latitudes where daylength is stably equal to 12:00 h has intrigued many authors. In rice, Dore (1959) showed that the duration to flowering of the variety Siam 29 varied between 161 and 329 d with sowing month in Malacca, Malaysia (2°12′N) where annual photoperiod variation was only 14 min. Under an artificial stable photoperiod from 11:50 to 12:10 (20 min range), Siam 29 showed a strong photoperiodic reaction but the duration to flowering only varied from 80 to 150 d, much less than that in the monthly sowing experiment. At a larger scale, Borchert *et al.* (2005) gathered a large dataset on the flowering time of 41 species of forest trees in Central and South America that exhibited synchronous bimodal flowering distribution in spring and autumn at equatorial latitudes, from 2.5°N to 5°S. In tropical zones at 10°N or 16°S, the flowering distribution was monomodal and synchronous with one of the equatorial flowering peaks. The authors pointed out that the bimodal distribution was synchronous with the peaks in the annual variation in the change in sunrise and sunset hours at the equator, in March-April and September-October.

Indeed, as a consequence of the ellipsoidal orbit of the earth around the sun and of the inclination of the axis of rotation of the earth, variations in the sunrise (SR, Fig. 1A) and sunset (SS, Fig. 1B) hours results in symmetrical variation in the daylength across the year (PP, Fig. 1C). However, SR and SS are asymmetrical across the year and so are their daily derivatives (dSS & dSR, Fig. 1D). All these vary with latitude. Similarly, Ramírez *et al.* (2010) reported that mango trees grown in La Mesa, Colombia, (4°31′N) flowered twice a year in separate sections of the canopy.

**Fig. 1:**
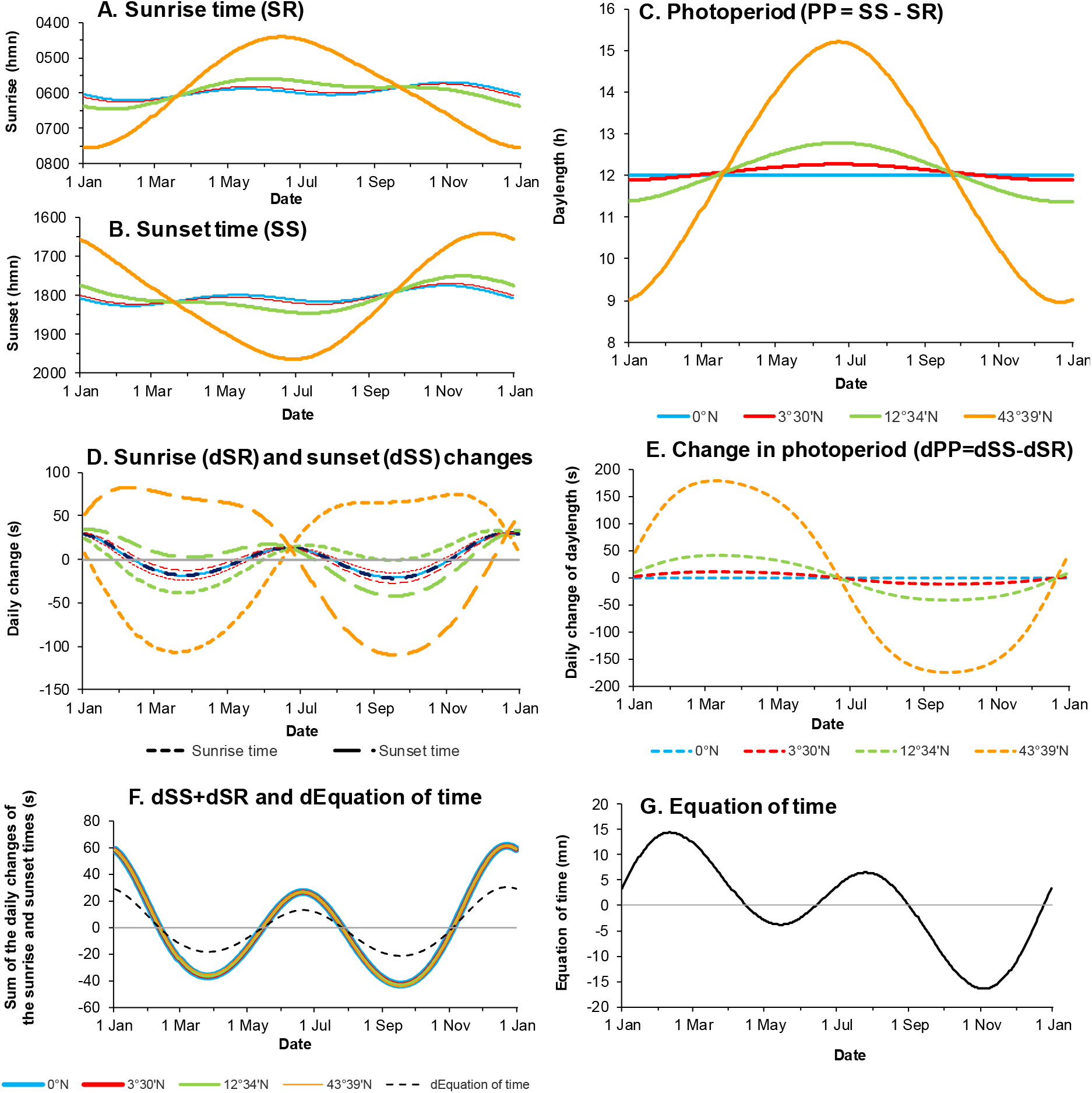
Daylength (A), sunrise time(B) and sunset time(C) with the year date and their respective derivatives, the daily rates of change (D, E) at 4 latitudes from the Equator to Montpellier, France. At any latitude, the sum of the daily rates of change of the sunrise and sunset times (F) was the double of the derivatives of the Equation of time(G).

In birds, the annual cycles of reproduction, moulting, and migration of high-latitude birds are rigidly controlled by seasonal variations in photoperiod (Gwinner and Scheuerlein, 1999). The stonechat birds living in Nakuru, Kenya, (0°14′S) initiate breeding at the onset of the rains in March or April (Dittami & Gwinner, 1985), whereas other populations of the same bird species living in Mount Meru, Tanzania (3°23′S) start breeding in October. Goymann *et al.* (2012) demonstrated that the reproductive cycle of stonechats living indoors in Germany with 12:00 h of stable day length and changes in the sunrise and sunset hours mimicking the changes in Nakuru was well synchronised among birds and with the cycle of wild birds living in Nakuru. Thus, they demonstrated the efficiency of the cue proposed by Borchert *et al.* (2005) (the daily change in sunrise and sunset times) in synchronising the reproductive phase. Since 2000, physiological studies have contributed much information on the relationships between the circadian clock and the seasonal triggering of the reproduction in plants (Shim and Jang, 2020). The circadian clock in each cell is made up of antagonist molecule couples that translate the day-night alternation into molecular concentration alternations. Three main genes encode three proteins (TOC1, CCA1, and LHY) that interact in a series of interlocking transcriptional feedback loops (Barak *et al.,* 2000). CCA1 and LHY concentrations peak at dawn, whereas that of TOC1 peaks at dusk. The action of the feedback loops is reversed at dawn and dusk, perceived through photoreceptors, including a blue photoreceptor, the cryptochrome, discovered in 1993 (Ahmad and Cashmore, 1993). Through a complex network of proteins with daily-oscillating concentrations, the soluble FT protein (Flowering LocusT) is synthetised in leaf cells at concentrations that are adjusted by both photoperiod and temperature (Kinmonth-Schultz *et al.,* 2018). The FT protein moves to the stem apex where it induces the transition to the reproductive phase.

The current paper relies on phenology data recorded during the last 26 years by sorghum scientists from the French Agricultural Research Centre for International Development (CIRAD) who conducted their research programmes in various locations from equatorial to temperate latitudes. Sorghum is a staple food in West-African Sahelian countries where CIRAD and ICRISAT developed many partnerships for sorghum crop improvement. It soon appeared that photoperiod-sensitivity was a key factor in the variety adaptation in this part of the world and thus deserved a better understanding. CIRAD also conducted sorghum breeding activities in Nicaragua with the International Center for Tropical Agriculture (CIAT) and the Nicaraguan Institute of Agricultural Technology (INTA) and in Colombia with the Colombian Corporation for Agricultural Research (CORPOICA). CIRAD and the International Rice Research Institute (IRRI) share a long tradition of partnership in research for rice crop improvement. Complementary monthly sowings of rice were thus conducted at IRRI headquarters in the Philippines, which helped to link results from contrasting latitudes. After a long maturation, an improved and simple model emerged that synthesised the current knowledge and could predict the response of sorghum and rice crops to photoperiod at any latitude. This model is described below.

## MATERIALS AND METHODS

### Data acquisition

#### Monthly sowings of sorghum in Mali

Four series of monthly sowings on approximately the 10^th^ day of each month were conducted from July 2000 to October 2008 at the Samanko ICRISAT research station near Bamako, Mali (12°34′N, 8°04′W, 330 m asl). Three tropical sorghum cultivars, CSM 335, IRAT 174, and Sariaso 10 were used during the first series (26-month study, Clerget *et al.* (2004, 2008)) and 12 new cultivars, plus CSM 335 as a control, were used during a second series (29-month study, Dingkuhn *et al.* (2008)). Seven East African cultivars, plus CSM 335, IRAT 174, and Sariaso 10 were used during a third series (12-month study). Finally, six cultivars from two distinct genetic clusters of durra sorghums (Deu *et al.*, 2006) were tested during the last series (15-to 18-month study) (Supplementary Table 1). Individual plots consisted of four 5 m-long rows, sown at 0.75 m × 0.20 m spacing. Split fertilisation doses were applied to secure the optimum growth of the plants (gypsum at 100 kg ha^−1^ and NPK at 128, 92, and 60 kg ha^−1^ as urea, diammonium phosphate, and KCl, respectively, during the first 45 d. Then, in late varieties N at 46 kg ha^−1^ was applied as urea monthly until the flag-leaf appearance). Soil moisture was never limiting because of biweekly irrigation during the dry season. Hand-weeding and insecticide were applied when required. Ten plants of the two central rows per plot were specifically labelled and the numbers of leaves that emerged from the whorl on the main culm were recorded weekly. From seedling emergence to PI, two plants per variety were sampled every week and after a longer interval for very late varieties. The plants were dissected to count the number of leaves initiated at the apex and to record the PI date.

Additionally, a sorghum core collection of 210 accessions established by CIRAD and ICRISAT scientists (Grenier *et al.,* 2001) was sown in Samanko, Mali on 13 November 2007 in an irrigated field. Plots consisted of one 3 m-long line distributed in an unbalanced incomplete block design, including six repeated checks. Adequate fertilisation and irrigation were applied to secure optimum growth, as described above. Dates at which 50% of the plants had reached flag-leaf expansion, heading, and flowering were recorded in each plot.

#### Simultaneous sorghum sowings in France and Mali

In France, sorghum experiments were managed by the CIRAD sorghum breeding program for the adaptation of tropical resources to temperate environments in Montpellier, France. Sorghum trials were sown in the GEVES Lavalette Research Station (43°39′N, 3°52′E) from mid-May. After ploughing, 300 kg/ha of fertiliser compound (N-P-K: 0-23-19) was applied, sowing was conducted at 75 × 20 cm, and sprinkler-irrigation was applied every 2 weeks when required. Three N-fertigations at 40 kg ha^−1^ N each were applied at approximately 30, 60 and 90 days after sowing.

From 1994 to 1997, two photoperiod-sensitive sorghum varieties (SSM 973, IS 9508) and one photoperiod-insensitive variety (IS 2807) were sown both in Samanko, Mali (17 Jun, 1, 15, and 29 Jul 1994; 23 Jun, 1, 9, and 20 Jul 1995; 16 Jun, 1, and 22 Jul 1996) and in Montpellier, France (18 May and 8 Jun 1994; 15 May and 17 Jun 1997), and the dates of flag-leaf appearance and flowering were recorded.

In 2001, CSM 335, IRAT 174, and Sariaso 10 were simultaneously sown in Samanko and Montpellier on 11 May, 7 June, and 3 July. Leaf appearance kinetics and PI were recorded as described above for monthly sowings experiments in Mali.

In 2006, five sorghum varieties (CSM 335, IRAT 174, IRAT 204, Sariaso 10, and Souroukoukou) were simultaneously sown in Samanko, Mali, and Montpellier on 19 May, 22 June, and 20 July. Leaf appearance and PI were also recorded and all 111 remaining plants of the late varieties were dissected on 17–18 October 2006, before the beginning of the cold season, to determine their reproductive stage. From 2008 to 2017, additional observations of flowering dates were conducted in Montpellier for the varieties Sariaso 10 and Sariaso 14 with sowing dates at approximately mid-May.

#### Sorghum in Nicaragua and Colombia

From 2002 to 2008, CIRAD conducted a regional participatory sorghum breeding program based in Nicaragua, in partnership with the Nicaraguan Institute of Agricultural Technology (INTA), the International Center for Tropical Agriculture (CIAT), national NGOs, and local farmer organisations, targeting resource-poor farmers in drought-prone areas where sorghum is an important staple crop. Local cultivars from Nicaragua, known as criollo or maicillón, generally grown in intercropping systems with maize and/or beans, are very late and highly photoperiod-sensitive and the quantification of this trait was included in the research program. African varieties (Sariaso 14 and Souroukoukou) were included in these experiments.

From 2008 to 2011, CIRAD and the Colombian Corporation for Agricultural Research (CORPOICA, now AGROSAVIA) implemented a research project to develop sweet sorghum hybrids adapted to four targeted agroecologies of Colombia. In the framework of this project, phenology, as well as other adaptive and productivity traits of a large collection of diverse cultivars and inbred lines, were recorded at three low latitude locations (Palmira, 3°30′N; Espinal, 4°26′N; and Monteria, 8°44′N). A large database was thus established from which data for some cultivars grown both in Nicaragua and Colombia with multiple sowing dates were selected.

#### Monthly sowings of rice in the Philippines

Published results of monthly rice sowings in Malaysia, close to the equator (Dore, 1959), and in Japan, at a temperate latitude (Kawakata and Yajima, 1995), constituted a good basis for comparison with monthly sowings conducted in the tropics. The six varieties from the Japanese experiment were obtained from the IRRI’s Gene Bank and rejuvenated in four pots per variety, which were sown in December 2011. Plants were then grown in a greenhouse at the IRRI farm, Los Baños, Philippines (14°11′N, 121°15′E) using a protocol as similar as possible to the original protocol. On or close to the 10^th^ of each month, 12 sowings were conducted during 12 consecutive months, starting on 10 May 2012. For each variety, three plants were grown in two 13 L-pots filled with soil from the IRRI upland farm. Fertilisers were applied in each pot (0.16 g P as SSP, 0.16 g K as KCl, and 0.06 g ZnSO4 at sowing; 0.16 g N as urea at 18 and 30 d after sowing and then monthly). Soil water content was kept at the saturation point for 2 weeks; then, the water level was maintained ~3 cm above the soil surface. Leaf appearance was recorded for the main stem of each of the six plants until the flag-leaf expanded.

The second series of 15 consecutive monthly sowings from 14 March 2014 only involved the variety Siam 29, used by Dore (1959). Seeds from two accessions, IRGC 13741 and IRGC 27, were obtained from the IRRI’s Gene Bank, but IRGC 27 was later discarded because of the earliness of the plants. The first sowing was conducted in six 13 L-pots placed in the IRRI greenhouse as in the series described above. All the following sowings were conducted in a small 6 × 6 m IRRI screenhouse, i.e. a field space covered with a 2 m-high metallic structure covered with a metallic mosquito net. Each month, six plants were sown at 20 × 20 cm in a 1 ×1 m bonded plot and managed as in the previous series. Leaf appearance was recorded on the main stem of each of the six plants until the flag-leaf expanded. In December 2015 and January 2016, simultaneous experiments were conducted in the greenhouse and the screenhouse to check for a possible effect of the enclosures.

### Data analysis

#### Duration from sowing to PI

The duration of the vegetative phase until the induction of flowering varies with photoperiod and temperature during the vegetative phase. Thus, the duration to flowering induction, i.e. PI in Poaceae, must be either known or carefully estimated before it can be related to the factors responsible for its variation. PI was recorded in all sorghum experiments conducted in Samanko where both leaf initiation and leaf appearance kinetics were recorded. In rice, the leaf appearance kinetics were recorded in all experiments conducted in Los Baños and the data were sufficient to accurately estimate the time of PI. For data from other sources, only the duration to heading or flowering was recorded. The duration to PI was then estimated with bilinear relationships parametrised from the large Bamako and Los Baños databases, as described below.

#### Ex-post estimation of PI date from the leaf appearance kinetics in rice

In the monthly sowings of rice conducted in Los Baños, only the flowering dates were recorded because the *ex-post* estimation of the PI date from the leaf appearance kinetics was easy and highly reliable in this species. In this specific species, a stable, re-checked, number of four developing leaves is hidden inside the leaf sheaths at any time during the vegetative phase until the time of PI (Yoshida, 1981).

The first step was to determine the parameters of leaf appearance kinetics in each variety. Depending on the variety and the sowing date, the dynamics of leaf emergence were either linear, bilinear, or trilinear and were fitted with linear or broken-linear segmented models, as previously detailed (Clerget *et al.,* 2008). In short, observed leaf number (LN) (initiated, emerged, ligulated, or senesced) was regressed against the elapsed thermal time from emergence (TT) using the following equations in the case of trilinear kinetics:

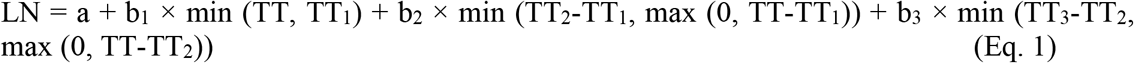

where a is the number of leaves at emergence, b_1_, b_2_, and b_3_ are the first, second, and third rates of development, and TT_1_, TT_2_, and TT_3_ are the thermal times when either the change in the rate occurred or the development terminated. Parameters were iteratively estimated using the procedure NLIN in SAS (2012). Phyllochrons were calculated as the reciprocal of the slope coefficients for leaf emergence.

Then, when for instance the leaf appearance was trilinear, the thermal time at PI (TT_PI_) was estimated as the thermal time at the appearance of the leaf (N-4), with N being the total number of leaves produced by the main stem:

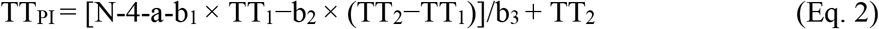

#### Relationship between the duration to flowering and the duration to PI

Bilinear relationships between either the observed or estimated duration to PI and the duration to flowering were fitted for sorghum in Samanko and rice in Los Baños data from the monthly sowings, as:

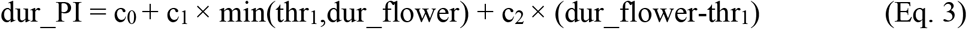

where dur_PI is the duration from sowing to PI, dur_flower is the duration to flowering, c_0_, c_1_, c_2_, and thr_1_ are parameters (Fig. 1 and Table 1). This relationship was used to estimate the date of PI from the flowering date in datasets in which PI was not recorded.

**Table 1:** Parameters and statistics of the bilinear relationship duration to PI = f(duration to flowering) plotted in Supplementary Fig. 1.

| Species | $c_0$ | $c_1$ | thr1 | $c_2$ | n | $R^2$ | RMSD |
| --- | --- | --- | --- | --- | --- | --- | --- |
| Sorghum | -12.68 | 0.689 | 135.8 | 0.892 | 859 | 0.969 | 8.52 |
| Rice | -9.54 | 0.569 | 111 | 1.024 | 94 | 0.9877 | 3.36 |

#### Using the sunrise and sunset times in modelling the date of PI at any latitude

The more recent models for the duration to flowering in response to the photoperiod used photoperiod thresholds that determined the onset and the end of the period during which flowering initiation was inhibited in qualitative photoperiodic varieties (Carberry *et al.,* 1992; Folliard *et al.,* 2004; Dingkuhn *et al.,* 2008). Therefore, several attempts were conducted to replace the photoperiod thresholds by (dSR + dSS) thresholds that would be similar at every latitude. However, the simpler additive model using PP, dSR, and dSS fit the observations better than any of the other more sophisticated models and was thus retained.

The 2.0 photoperiod model was built based on the model proposed by Major (1980). Seedling emergence is followed by a juvenile phase, known as the basic vegetative phase (BVP). From the end of the BVP, begins the photoperiodic sensitive phase (PSP) of genetically determined duration that ends at PI. Under stable photoperiod and controlled environments, PI occurs immediately after the end of the BVP when the daylength (PP) is below the base photoperiod (Pb_1_). Otherwise, when PP exceeds Pb_1_, the duration of PSP increases linearly with PP as a product of a photoperiod-sensitivity coefficient (Ps) with the difference between the actual photoperiod and Pb_1_ being (Ps×(PP-Pb_1_)) in varieties with a quantitative short-day response.

In natural environments, the expected duration of PSP on a specific day PSP(day) = f(PP, T) changes every day with PP and T (the mean daily temperature); thus, it was assumed that plants accumulate daily progress towards PI, according to the equation dPI = 1/f(PP, T) (Summerfield *et al.* 1992). PI occurs when the daily sum of progress reaches 1.

The daily change in sunrise (dSR) and sunset (dSS) times is a sufficient cue to trigger flowering in trees and to synchronise the circannual reproductive rhythms of birds living at the equator at similar dates as in congeners living in the tropical areas (Borchert *et al.,* 2005; Goymann *et al.,* 2012). Thus, an additive effect of each of these two factors (dSR and dSS) was introduced in the crop-photoperiodism model 2.0, by multiplying them by the respective sensitivity coefficients, SRs_1_ and SSs_1_. In fields, these two effects act on any day of the year, whereas the PP effect acts only when PP > Pb_1_. However, it appeared that in rice grown in the greenhouse all three effects act only when PP > Pb_1_. When the linear combination of the three effects falls to one or is lower than one, 1/PSP(day) = 1 and PI occurs on the same day.

This model could fit the data of the rice variety Siam 29 when grown close to the equator with June to December sowings but not with January to May sowings. During the latter time interval, the duration to PI decreases by one month between each monthly sowing date, indicating the absence of any progress towards PI. The modelling solution was to hypothesise that PSP(day) was very long from approximately the spring equinox to the summer solstice when both the photoperiod (PP) is above a specific threshold PPb_2_ and dSS > dSR because of a stronger response to dSR and dSS with larger coefficients SRs_2_ and SSs_2_. The concept was then reused in modelling the response of IRAT 204 with sowings from July to September, the time when PP > PPb_2_ and dSS < dSS, and by extension to improve the model fit of nearly every sorghum and rice variety for sowings from June to August.

The time unit was changed from thermal time to day based on further detailed results and discussion. In summary, the PI of plants simultaneously sown in Bamako (Mali) and Montpellier (France) occurred on close dates but after a longer thermal time for plants grown in Mali. The duration in days to PI of the daylength-insensitive variety IRAT 204 was also insensitive to the mean temperature in Bamako as in Montpellier. These results were attributed to acclimation of the duration to PI to the temperature at seedling emergence, which was able to buffer the effect of temperature on the plant development rate within a large range of temperatures. Given the current knowledge, it thus appeared more accurate to assume the insensitivity of the duration to PI in days to the temperature in usual cropping conditions than the contrary, even if not entirely true.

Finally, the 2.0 model can be summarised as below:

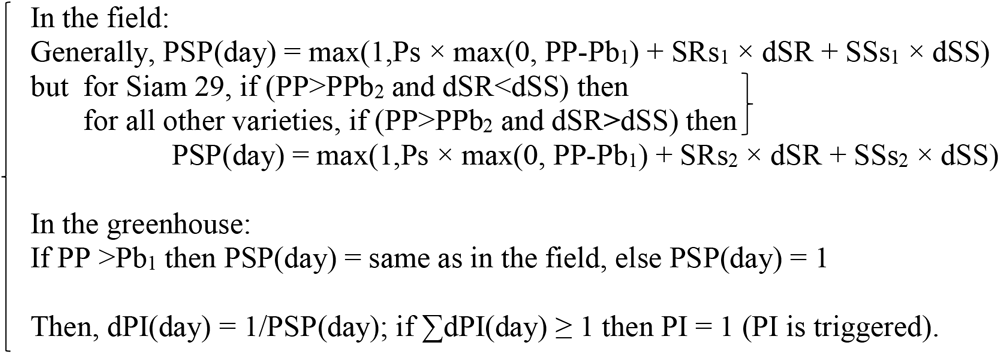

Daily photoperiod and sunrise and sunset times were calculated using the equations from Laureanno (1992) (Supplementary Excel file). Parameters for the model were issued from an iterative optimisation using the evolutionary solving method in the Microsoft Excel Solver to minimise the sum of square errors (predicted-observed). When data were available from several latitudes, data from one to two latitudes were used to calibrate the parameters of the model that was then run for validation at the other latitude.

The accuracy of the prediction of the duration to PI was assessed using the coefficient of determination (*R*²) derived from the regression of observed and predicted days to PI, and the root mean square deviation (RMSD), which represents a mean weighted difference between predicted and observed data. RMSD was calculated as RMSD = [(observed-predicted)²/n]^0.5^. The computations were conducted for all sowing dates and on a dataset reduced to eight sowing months, from March to October, to discard the highly contrasting durations to PI observed from November to February sowings in some of the varieties. Prediction of the duration to PI for daily sowing dates was then computed using the model with the optimised parameters in SAS data steps. Proc GLM (SAS, 2012) was used to compute the ANOVAs.

## RESULTS

### Universalsolar cue fromht e Equator to the Poalar Circles

Daylength and its derivative, the daily change in daylength, have symmetrical yearly variations centred on the equinoxes (21 June and 21 December) (Fig. 1C & E). The daily change in daylength is also the difference, dSS-dSR, whereas the sum of the daily changes in the sunrise and sunset hours, dSS + dSR, is always the same everywhere on earth from the Equator to the Polar Circles (Fig. 1F). This sum of dSS+dSS is equal to the double of the daily derivative of the equation of time, i.e. of the difference between the solar time (indicated by sundials) and the mean solar time (shown by clocks) (Fig. 1G). However, thresholds of this cue were unsuccessfully tested to replace the previous thresholds of the photoperiod in qualitative photoperiod-sensitive varieties.

### Durations of the vegetative phase in monthly sowings in the tropics

#### Monthly sowings of sorghum in Samanko, Mali

The 28 sorghum varieties that were sown monthly in Samanko, Mali from 2000 to 2008, showed a broad range of responses to the sowing date (Fig. 2, Fig. 3A-C, and Supplementary Fig. 2). Results for the 14 varieties sown between 2000 and 2005 have been previously reported (Clerget *et al.,* 2004; Dingkuhn *et al,.* 2008). Another set of 14 varieties was sown from 2005 to 2008. Eight varieties were selected that covered the entire range of observed patterns of photoperiodic responses of the duration to PI to the sowing date, from insensitive to highly sensitive varieties (Fig. 2). IRAT 204 was insensitive during eight sowing months, from November to June, but durations to PI were longer from July to October (Fig. 2A). CSM 63-E was insensitive in sowings from May to October and showed longer durations to PI for sowings from November to April (Fig. 2B). Fig. 2C to G show varieties with increasing photoperiod-sensitivity for sowings from January to September. Sima and Ouéni were quantitative photoperiod-sensitive, whereas IRAT 174, Kaura D-12, and IS 15401, with a breakpoint in the annual pattern between two early sowing dates in the year, showed qualitative photoperiodism. The shorter duration to PI occurred for the September and October sowings in every variety. The duration to PI was much longer for November and December sowings in the varieties CSM 63-E, IRAT 174, and Ouéni.

**Fig. 2:**
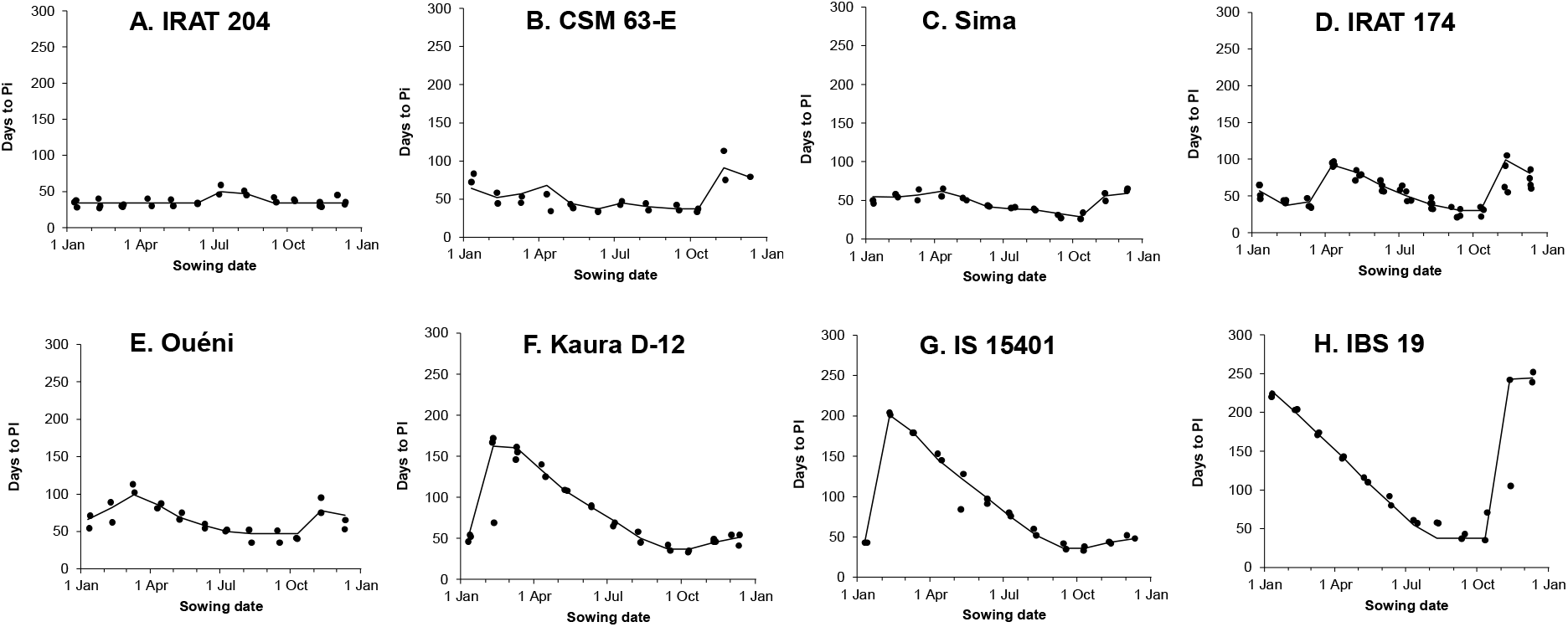
Observed duration to panicle initiation in two-year monthly sowing series (●) of eight sorghum varieties covering the whole observed range of photoperiod-sensitivity grown in field at Samanko (12°34’N) and their estimation by the crop- photoperidism model 2.0 (─).

**Fig. 3:**
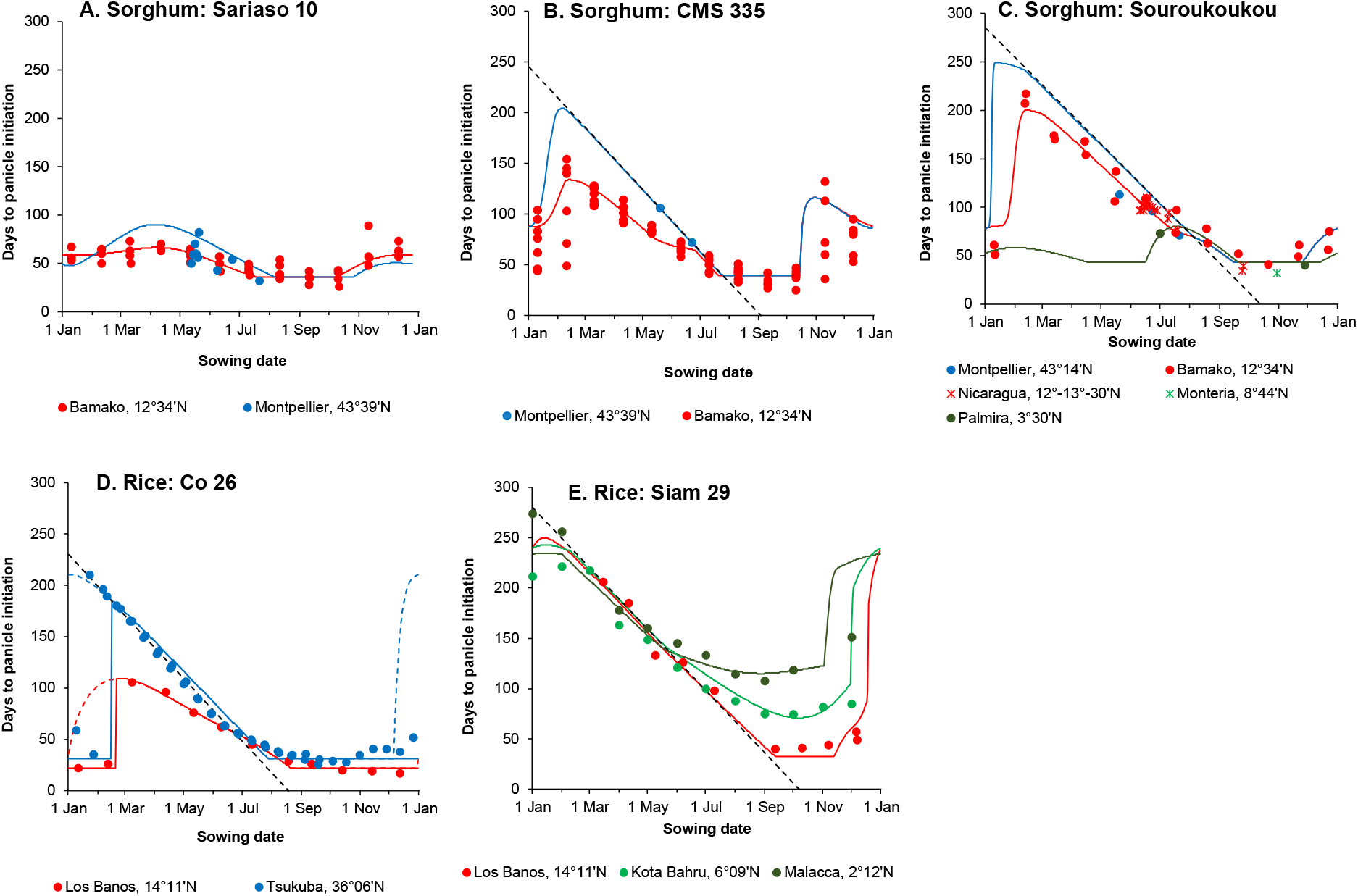
Observed durations to panicle initiation (●) and daily model prediction (─) for three sorghum varieties monthly sown in Samanko (Mali) and additionally observed on the usual planting date at either one other latitudes (AB&) or four other latitudes (C);and for two late rice varieties monthly sown at either two (D) or three (E) latitudes. The −1:1 line was drawn in late varieties (---).

A new pattern was observed for the first time in four landraces from South Tanzania (IBS 19, 30, 40, and 582) (Fig. 2H and Supplementary Fig. 2O-Q). In these highly photoperiod-sensitive varieties, the PI of the November and December sowings could be delayed until July or August of the next year though with a large inter-annual variation. PI of all sowings from January to July occurred during August. This November threshold in the photoperiodic reaction was confirmed by the duration to flag-leaf observed for 207 accessions of the sorghum core collection sown at Samanko on 13 November 2007, which ranged from 59 to 350 d. Sixteen accessions of this diversity panel reached PI after 1 September 2008, i.e. 9.5 to 12 months after the sowing date (Supplementary Table 2).

#### Monthly rice sowings in Los Baños, Philippines

A series of 12 sowings in the greenhouse replicating the experiment of Kawakata and Yajima (1995) was fully successful. The five Japanese varieties and IR 72 sown monthly in Los Baños showed only small changes in the duration of the vegetative phase in response to the sowing date, with a maximum duration for the May sowing and minimum for the September to February sowings (Fig. 4). Conversely, the duration to PI of the Indian variety Co 26 strongly reacted to the sowing date with a maximum duration for the March sowing and minimum for the December sowing (Fig.3D).

**Fig. 4:**
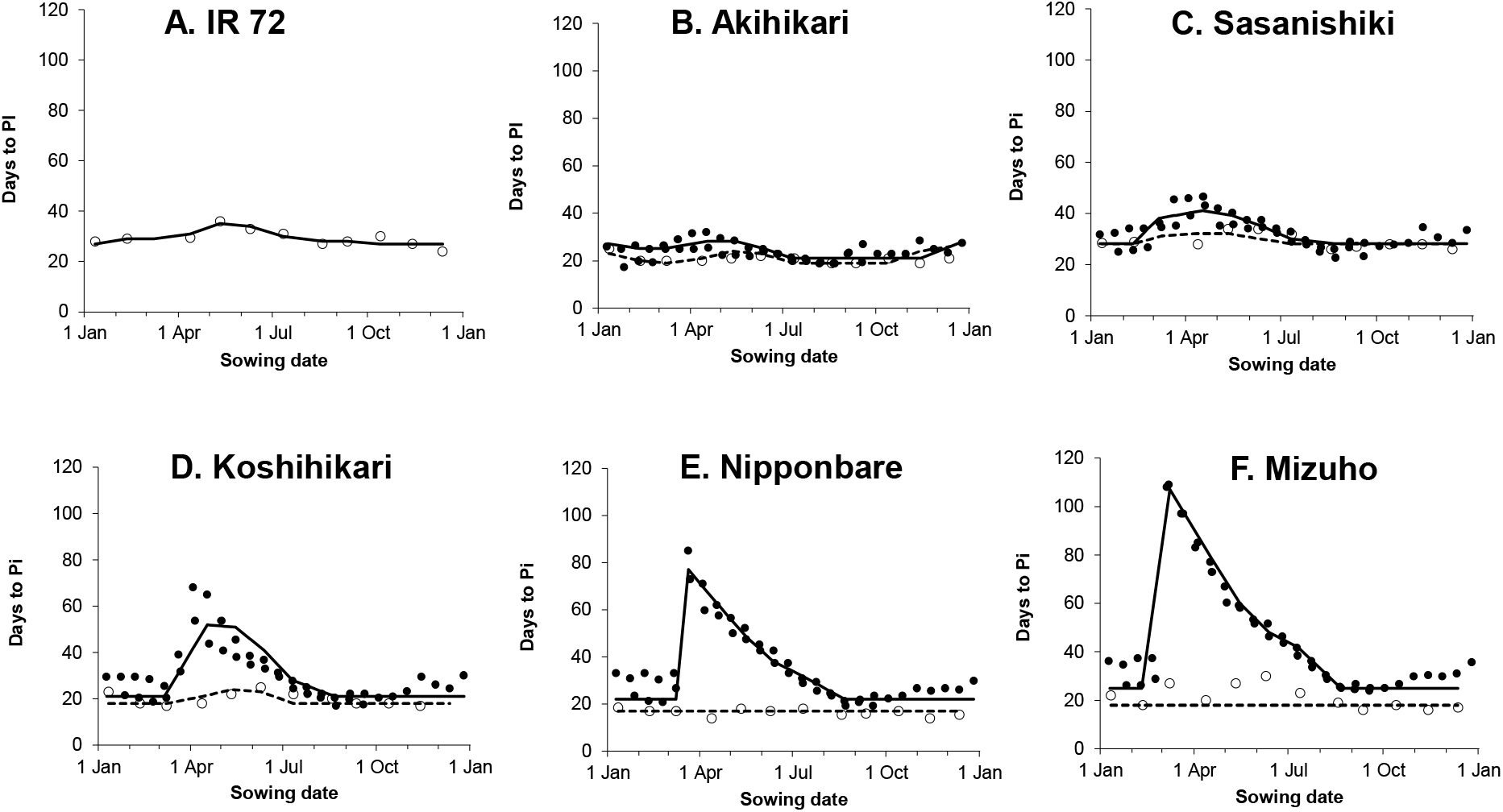
Duration to panicle initiation during a 2-year fortnightly sowing series (●) of five rice varieties grown in a greenhouse in Tsukuba (36°06’N) and during a 1-year monthly sowing series (○) of six rice varieties grown in a greenhouse in Los Baños (14°11’N). The model calibrated with Tsukuba data (─) was validated with photoperiod data at 14°11’N **(…).**

Only the first eight monthly sowings of Siam 29 in the series replicating the experiment of Dore (1959), from March to December 2014, were successful. None of the following sowings could be maintained until PI because of stem borers damage to the January and February sowings. Then, Tungro viruses infestation damaged all the plants from the next sowings. The time to PI was at a maximum for the March sowing and decreased for the next sowings until reaching its minimum for the September to December sowings (Fig. 3E).

### Comparison of the duration to PI from the Equator to the temperate latitudes

#### Simultaneous dates of PI from the equatorial belt to the temperate area for sorghum

Results for photoperiod-insensitive and intermediate-sensitive sorghum varieties strongly questioned the use of the thermal time to report the duration of the vegetative phase. IRAT 204 is reputed as being non-photoperiod sensitive but showed a response in the duration of the vegetative phase for the sowing dates from July to October in Bamako (Fig. 5A). For the eight remaining sowing dates, the duration of the vegetative phase was stable (mean = 33.4 d) and insensitive to the latitude, and consequently, to both the day length and temperature when sown in mid-May in Montpellier where mean temperatures are always lower than those in Bamako (Fig. 5B) (mean = 35 d). For the eight sowings from November to June, the slope of the regression between the duration to PI and the mean temperature did not significantly differ from zero (−0.22 d °C^−1^, *P* = 0.36), within a 19–31.5 °C range of mean temperature (Fig. 5C). PI of the moderately photoperiodic sorghum varieties Sariaso 10, Sariaso 14, IS 2807, and SSM 973 occurred on the first fortnight in July for May sowings and in late-July to early-August for June sowings, independent from the latitude of the location, either 12–14°N or 43°39′N (Fig. 6 A-D). In such situations, the thermal time was much shorter in Montpellier than in Bamako. Thus, in 2001, Sariaso 10 was sown on 7 and 10 May in Montpellier and Bamako, respectively, and reached PI after 50 d or 502°Cd and mean daylength of 15 h in Montpellier and after 54 d or 904°Cd and mean daylength of 13:35 h in Bamako (Supplementary Table 3). Phyllochron1, set at seedling emergence, was 35°Cd in Montpellier and 47°Cd in Bamako and similar differences were observed for June sowings and the varieties CSM 335 and IRAT 174. However, in 2006, tropical varieties flowered exceptionally late in both locations and PI occurred at similar thermal times. Sariaso 10 was simultaneously sown on 20 May 2006 in both locations and reached PI after 82 d or 984°Cd in Montpellier and 60 d or 978°Cd in Bamako. In 2006, phyllochron1 was later in Montpellier (41°Cd) than in Bamako (37°Cd). In Montpellier, temperatures were exceptionally high (up to 34°C) during the last week of May 2006 and again during the last dekad of June 2006 (up to 36.8°C) for the times of seedling emergence for May and June sowings. For May and June sowings, PI was delayed from the average by 1 month and 1 fortnight, respectively. Location and year had significant effects on phyllochron1, thermal time to PI, and the total number of leaves.

**Fig. 5:**
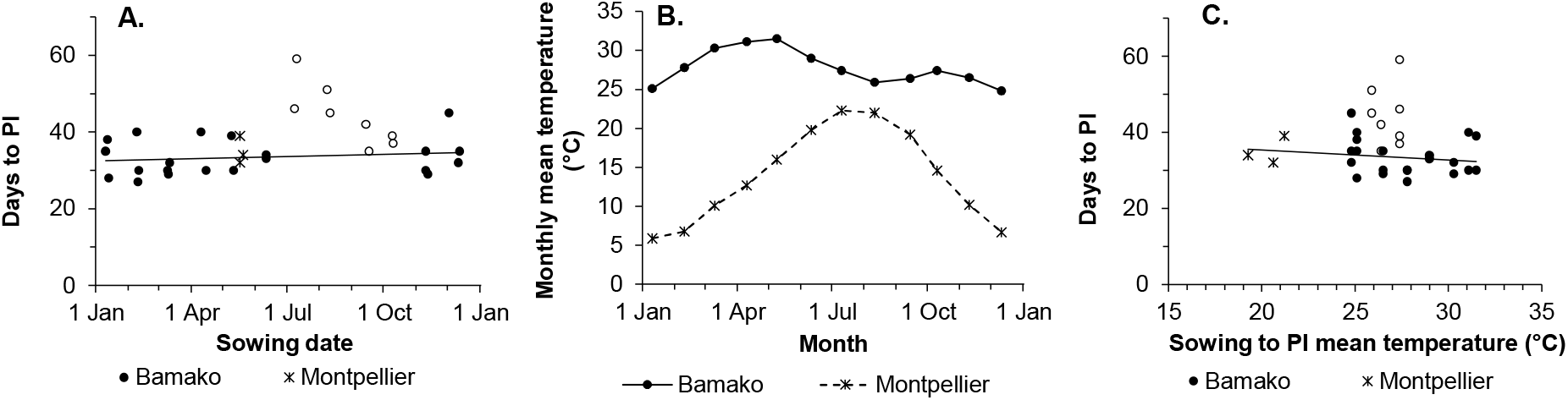
Observed durations to panicle initiation in days (A) by sowing date of the non-photoperiodic sorghum variety IRAT 204 monthly sown in Samanko, Mailfrom2003 to 2005 (● and ○) and in May 2001, 2010 and 2017 in Montpelier, France (*). In Bamako, a photoperiodic reaction was observed in sowings from July to October (○). B) Monthly mean temperatures in both locations and C) the relationship between the mean temperature and the observed durations to PI.

**Fig. 6:**
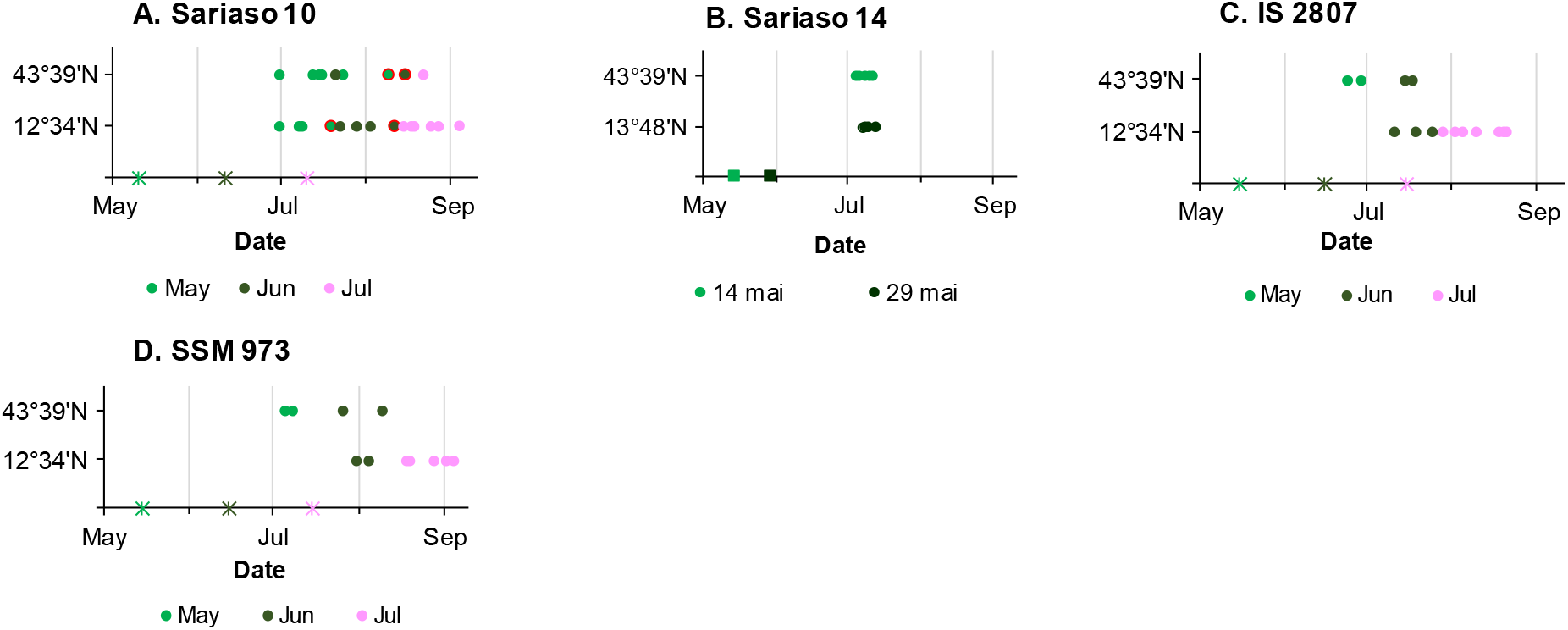
Distribution of the dates of panicle initiation of four sorghum varieties simultaneously sown in Montpelier (43°39’N) and either Bamako (12°34’N) or Somoto, Nicaragua (13°48’N) from May to July between 1994 and 2017. Divergent results from 2006 were marked with red circles around the symbol for Sariaso 10 (A).

The Malian landrace Souroukoukou was grown in six locations in Africa, Europe, and Central and South America, covering a range of latitudes from 3°30′N to 43°39′N and showed similar responses to the sowing date in the duration to PI at any latitude, including at low latitude in Palmira, Colombia (Fig. 3C). Six late sorghum varieties from Nicaragua grown at four latitudes from 3°30′N to 13°N showed the same independence to latitude in the response of the duration to PI from the sowing date as Souroukoukou, especially for the varieties 99 PREIME 117 and EIME 119 (Supplementary Fig. 3). Indeed, the duration to PI was shorter in Palmira for four varieties sown during late-June than that in the other locations.

#### Divergent responses to latitude in the duration to PI of the rice varieties

The IRRI’s bred variety IR 72 and the five Japanese varieties chosen for their contrasting photoperiod-sensitivity in Japan were insensitive to the sowing date when grown in a greenhouse at the tropical latitude of Los Baños (Fig. 4). Conversely, the Indian Co 26 tropical rice variety was highly sensitive to the sowing date at both temperate and tropical latitudes (Fig. 3D). The estimated dates to PI of Co 26 were earlier in the Philippines (14°11′N) than in Japan (36°06′N) for March to May sowing dates and was similar during other sowings.

The Malayan rice landrace Siam 29, known for its high sensitivity to the sowing date at latitudes close to the equator, showed an even stronger response to the sowing date when sown in the field at the tropical latitude of Los Baños (Fig. 3E). The PI dates of Siam 29 were late and synchronous in Malaysia and the Philippines for sowings from March to June–July. For sowings from July–August to December, the sowings in Los Baños confirmed that PI occurred earlier at higher latitudes where the photoperiod is shorter during this period in the year.

### Crop-photoperiodism model 2.0 fitted any type of photoperiodic response at any latitude

#### Monthly sorghum sowings in Bamako

For all sorghum varieties, a set of optimised parameters permitted the proposed model to well fit data at every sowing date, which was not feasible with the previous models for photoperiodism (Fig. 2, Fig. 3A-C, Supplementary Fig. 2, Table 2, and Supplementary Table 4). In particular, the parametrised model was notably well fitted to the longer duration to PI for sowings in November and December regardless of the amplitude of the increase without any temperature input. The November breakpoint in varieties, such as IBS 19 and the specific pattern of IRAT 204 from July to October, was also well fitted. The respective role of daylength and the changes in the sunset and sunrise times in the daily-predicted duration to PI were plotted for three sorghum varieties grown at several latitudes (Fig. 7A-G). The day length often played the main role that was modulated by dSR and dSS, although the respective weights of the components could be balanced, as in CSM 335 in Bamako. The Ps coefficient, expressed in d h^−1^, had much higher values than the SRs and SSs coefficients, expressed in d²s^−1^ because the choice of the units ensured that the different products in the linear model had homogeneous orders of magnitude.

**Table 2:** Parameters of the crop-photoperidism model 2.0 and data fitting of A) 8 sorghum varieties representative of the range of photoperiod-sensitivity observed in Bamako, of B) two sorghumvarieties grown in Bamako and in Montpellier, and C) of one late sorghum variety grown in Bamako, Montpelier and Cali. When data from several locations were available, they were used for either calibranting (C) or validating (Val) the model Data fitting was computed either for all data from twelve sowing months or for data from only eight sowing months, from March to October.

| Variety | Location | Parm | Parameters |  |  |  |  |  |  |  | Data fitting |  |  |  |
| --- | --- | --- | --- | --- | --- | --- | --- | --- | --- | --- | --- | --- | --- | --- |
|  |  |  | BVP<br>days | P<sub>b1</sub><br>h | P<sub>s</sub><br>day h<sup>-1</sup> | SR<sub>s1</sub><br>day<sup>2</sup> s<sup>-1</sup> | SS<sub>s1</sub><br>day<sup>2</sup> s<sup>-1</sup> | P<sub>b2</sub><br>h | SR<sub>s2</sub><br>day<sup>2</sup> s<sup>-1</sup> | SS<sub>s2</sub><br>day<sup>2</sup> s<sup>-1</sup> | 12 months<br>R<sup>2</sup> | RMSD | March-October<br>R<sup>2</sup> | RMSD |
| A. One latitude |  |  |  |  |  |  |  |  |  |  |  |  |  |  |
| IRAT 204 | Bamako | C | 33.04 | - | - | - | - | 12.02 | -0.056 | -0.498 | 0.58 | 4.8 | 0.69 | 4.6 |
| CSM 63E | Bamako | C | 35.99 | - | - | -0.2787 | 1.8779 | 12.12 | -0.5603 | -0.4638 | 0.70 | 11.2 | 0.29 | 10.0 |
| Sima | Bamako | C | 27.87 | 9.59 | 7.90 | -0.097 | 0.5194 | 11.44 | -0.653 | 0.2831 | 0.85 | 4.5 | 0.88 | 4.0 |
| IRAT 174 | Bamako | C | 19.0 | 10.83 | 28.62 | 0.7825 | 2.3817 | 11.42 | -0.215 | 0.5169 | 0.79 | 10.9 | 0.92 | 6.2 |
| Ouéni | Bamako | C | 46.0 | 11.92 | 46.27 | -0.3063 | 1.0656 | 12.26 | -1.160 | 0.3974 | 0.82 | 8.8 | 0.94 | 7.4 |
| Kaura D12 | Bamako | C | 36.0 | 11.83 | 172.77 | -0.2534 | 0.5210 | 12.34 | -0.111 | 1.9927 | 0.86 | 18.1 | 0.99 | 5.5 |
| IS 15401 | Bamako | C | 35.11 | 11.89 | 286.64 | -0.1605 | 0.4183 | 12.22 | 4.8888 | 2.1078 | 0.98 | 8.7 | 1.00 | 10.6 |
| IBS 19 | Bamako | C | 36.79 | 11.0 | 126.68 | -3.0956 | 3.9852 | - | - | - | 0.85 | 29.0 | 0.98 | 7.7 |
| B. Two latitudes |  |  |  |  |  |  |  |  |  |  |  |  |  |  |
| CSM 335 | Bamako | C | 38.35 | 11.92 | 110.37 | -0.510 | 3.1132 | 12.35 | -3.543 | 0.5383 | 0.66 | 7.6 | 0.95 | 5.8 |
|  | Montpellier | Val | “ | “ | “ | “ | “ | “ | “ | “ | - | 0.65 | - | - |
| Sariaso 10 | Bamako | C | 34.87 | 10.78 | 11.45 | -0.164 | 0.5012 | 12.74 | -0.753 | 1.1656 | 0.48 | 5.9 | 0.70 | 5.0 |
|  | Montpellier | Val | “ | “ | “ | “ | “ | “ | “ | “ | 0.42 | 13.3 | - | - |
| C. Three latitudes |  |  |  |  |  |  |  |  |  |  |  |  |  |  |
| Souroukougou | Bamako | C | 42.17 | 12.3 | 915.14 | -0.9634 | 0.5737 | 9.94 | -1.286 | -0.7193 | 0.93 | 13.8 | 0.94 | 11.9 |
|  | Cali | C | “ | “ | “ | “ | “ | “ | “ | “ | - | 2.2 |  |  |
|  | Montpellier | Val | “ | “ | “ | “ | “ | “ | “ | “ | 0.97 | 21.7 |  |  |

**Fig. 7:**
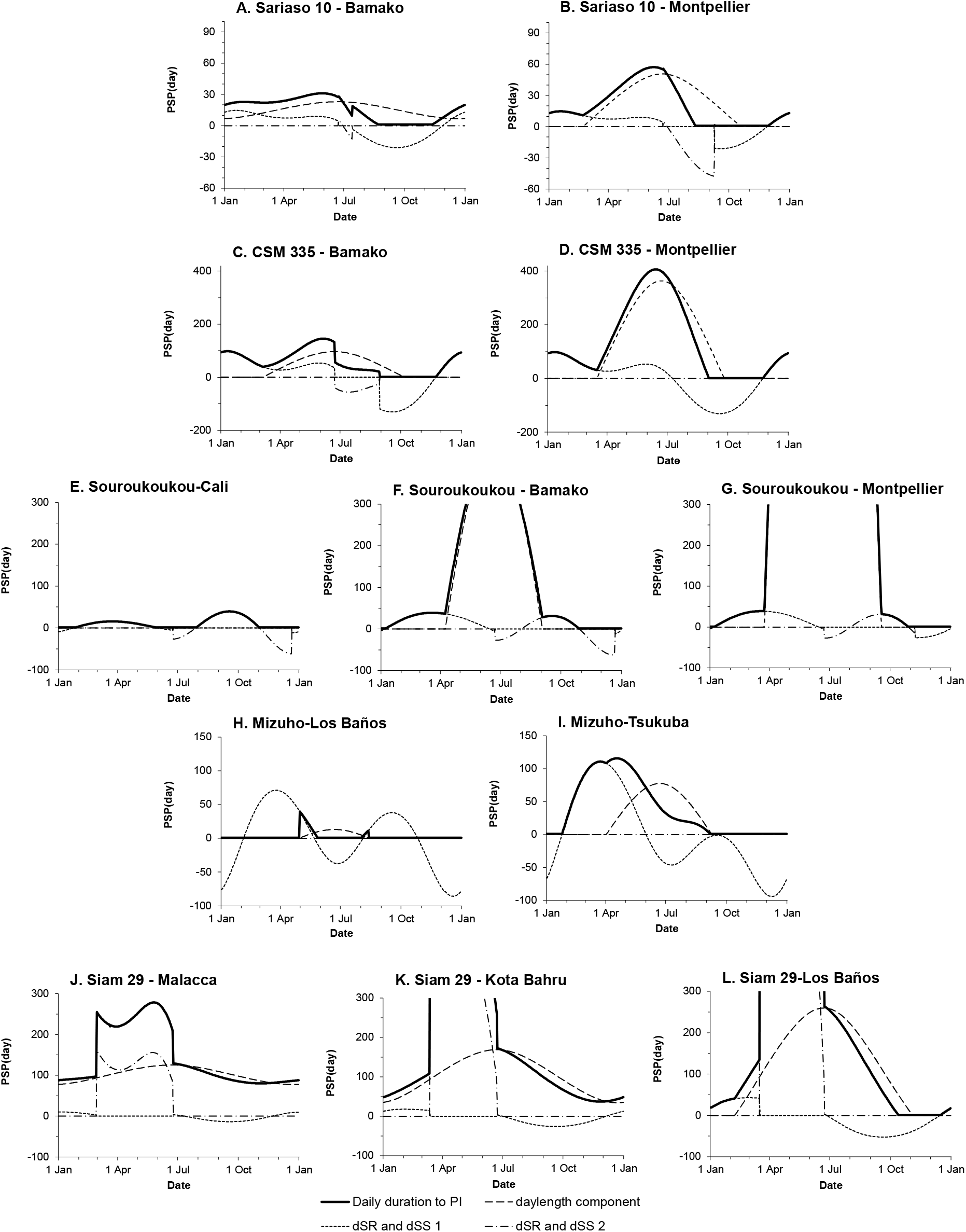
The daily-predicted duration of the PSP [PSP(day), in days] resulting from the addition of the daylength component and of the daily change in sunset and sunrise time component 1 or 2 for five varieties grown at several latitudes.

The predicted duration to PI of IRAT 204 and CSM 63-E did not respond to daylength (Ps and Pb were undetermined) but only to the additive effects of the sunrise and sunset changes (SRs and SSs coefficients in Table 2). As expected, the daylength parameter (Ps) was larger in varieties with a longer duration to PI with the noticeable exception of the IBS series. In the four IBS varieties, the SRs_1_ and SSs_1_ coefficients were much higher than in other varieties and stable over all sowing dates (no SRs_2_ and SSs_2_). BVP varied from 19 to 50 d and Pb_1_ from 8.2 to 12.3 h with no relationship with the level of photoperiod-sensitivity.

In many varieties, such as Kaura D-12 and IBS 19, the data fitting was better for the sowings from March to October than for data from the 12 sowing months because of high inter-annual variability in the duration to PI in some sowings from November to February (Fig. 2 and Table 2).

#### Sorghum varieties sown at several contrasting latitudes

Once calibrated with data from monthly sowings in Bamako, the model was run with data for the daylength and its changes in Montpellier, and the predictions were compared with the observations (Fig. 3 A-C and Table 2). For CSM 335 and Sariaso 10, RMSD values in Montpellier were higher than those in Bamako because the predictions of the duration to PI in Montpellier for May to July sowings were too long. Conversely, RMSD was very low for CSM 335 in Montpellier (two observations in 2006). Data from both Bamako and Palmira were needed to accurately calibrate the model for the variety Souroukoukou. Once done, the model well predicted the duration to PI in Montpellier for May to July sowings, and in Nicaragua, at similar latitudes as in Bamako. The duration to PI equal to the BVP was also accurately predicted for the October sowings in Monteria. However, the availability of only two sowing dates in Palmira were not enough to guarantee the validity of the yearly-predicted duration to PI, prospectively plotted.

#### Monthly rice sowing series at several contrasting latitudes

In rice, two long-time published monthly series in either Malaysia or Japan were complemented with two similar monthly sowing series in Los Baños, Philippines (Fig. 3D and E and Fig. 4). Thus, comparisons between monthly sowing series at several latitudes were available.

For the five Japanese varieties from the Tsukuba series, the response of the duration to PI to the sowing data was either null or very small in Los Baños (Fig. 4B to F) and the model was consequently calibrated with the data from Tsukuba. Conversely, the duration to PI of the Indian variety Co 26 responded strongly to sowing date in both locations and the model was calibrated with data from Los Baños because it better fitted the data from Tsukuba than did the reverse. No parameters were able to fit well with the observations for the five Japanese varieties. The highly variable duration to PI for January and February sowings of Co 26 grown in Tsukuba argued in favour of a breakpoint of the yearly pattern in early January, whereas the duration to PI of the other five varieties was minimal for sowings from September to February or March. The generality of this long low plateau in rice was questioned and found false. On the contrary, in the rice variety monthly sowings conducted in paddy fields in northern Senegal, the duration to PI and flowering was minimal for August– September sowings and longer for October to February sowings, similar to IR 64 or Nipponbare (Supplementary Fig. 4D and E) (Dingkuhn *et al.,* 1995, 2015). Unlike the Tsukuba series, observations from the Senegal series were well fitted to the crop-photoperiodism model 2.0 with no temperature input (Fig. 4 and Supplementary Table 5). It was thus hypothesised that the greenhouse modified the perception by the plants of the lighting cues for sowings from November to February or even March when the photoperiod was below Pb_1_. Under this additional constraint, the model fitted well the observations from Tsukuba. In Akihikari, Sananishiki, Co 26, and Koshihikari, Pb_1_ was estimated at 9.97, 12.0, 12.08, and 12.57 h, respectively, and the model predicted a response of the duration to PI to the sowing date in Los Baños where the maximum PP was 12.89 h (Fig. 4B-D and Fig. 3D). Conversely, for the two other Japanese varieties (Pb1 = 12.57 and 12.77 h), the model predicted the absence of a response to the sowing date, as observed and despite a strong dSR-dSS component similar to Mizuho (Fig. 7H).

In Siam 29, the crop-photoperiodism model 2.0 calibrated with data from Malacca and Kota Bahru, fitted well the monthly sowing series at the three contrasting latitudes (Fig. 3E and Table 3). The components dSR and dSS were hypothesised to cause the main lengthening of the predicted PSP, and thus, to play the main role in the very long durations to PI observed at low latitudes for sowings from January to June (Fig. 7J-K). At a tropical latitude, in Los Baños, the addition of the responses to PP, dSR, and dSS resulted in a very long-predicted PSP(day) (Fig. 7L).

**Table 3:**
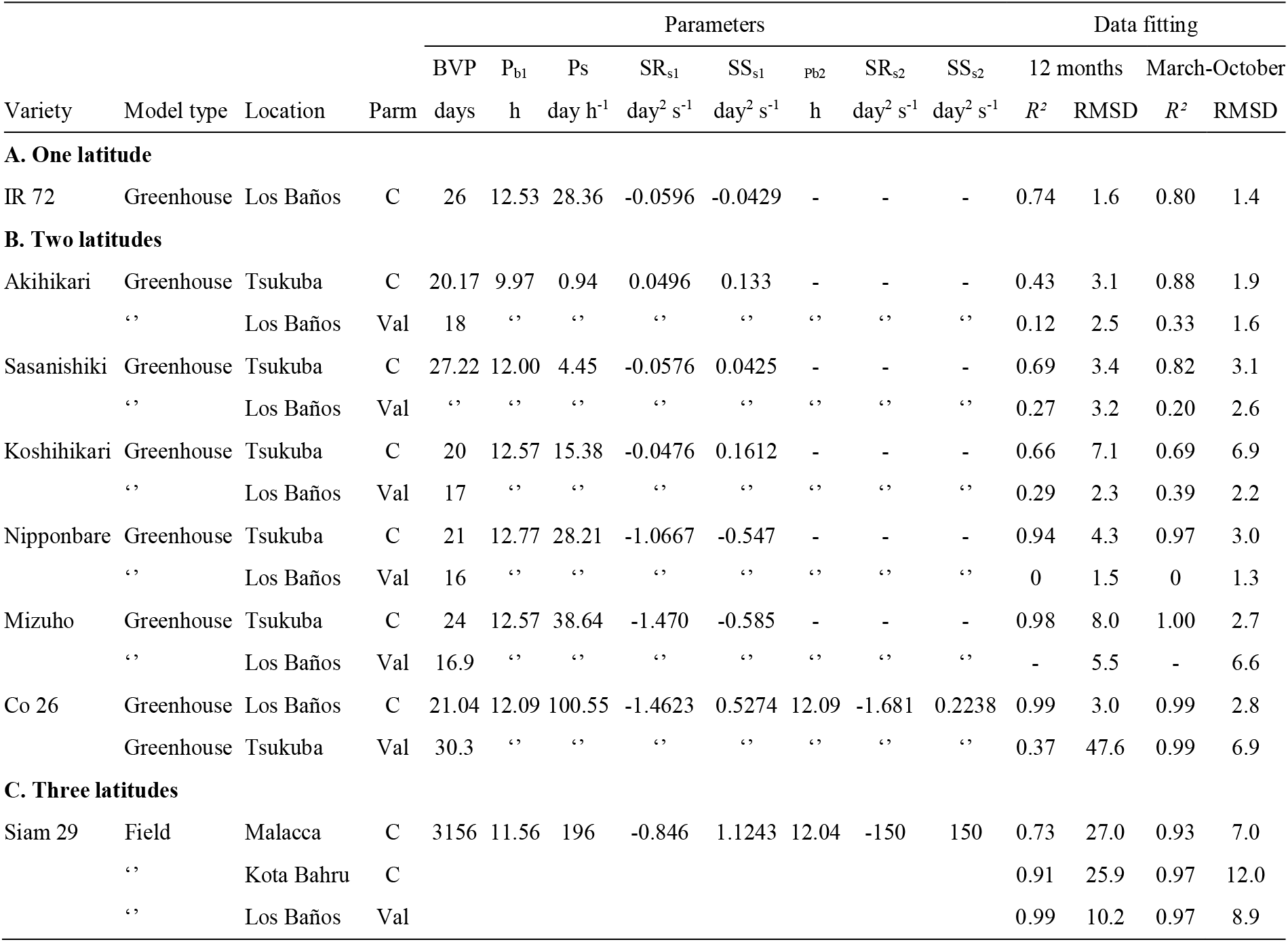
Parameters of the crop-photoperidism model 2.0 and data fitting of A) one rice variety grown in a greenhouse in Los Baños, of B) six rice varieties grown in greenhouse in Tsukuba and in Los Baños, and C) of one late rice variety grown in Malacca, in Koat Bahru and in Los Baños. When data from several locations were available, they were used for either calibrating (C) or validating (Val) the model. Data fitting was computed either for all data from twelve sowing months or for data from only eight sowing months, from March to October.

## DISCUSSION

### Duration to PI accmliaet s to the temperature

Durations are expressed in thermal time in the phenological models because temperature is a major factor in any plant metabolic activity, including the duration to flowering (Summerfield *et al.,* 1997). Many experiments conducted in growth chambers on various species strongly support this choice. However, photoperiod-sensitive varieties sown simultaneously in May and June in Bamako and Montpellier repeatedly flowered at close dates despite the highly contrasting photoperiod and temperature in both locations (Fig. 6). Such observations question both the photoperiod and the thermal time models. Indeed, May is a very hot month in Bamako (Tmean = 31.5°C), whereas germination and early growth can be difficult in Montpellier because of continued low temperatures (Tmean = 16°C) (Fig. 5B). Similarly, the duration to PI in days, limited to the BVP in the non-photoperiod-sensitive variety IRAT 204, was also insensitive to the temperature within a large range, 19.2 to 31.5°C, in contradiction with the current photothermal models (Fig. 5C). It appears that sorghum plants have the homeostatic ability to maintain the duration to PI at a rather stable number of days within a large range of mean temperatures by adjusting the thermal time duration. Kirby *et al.* (1985) previously hypothesised that in wheat and barley, the phyllochron was set in response to day length and temperature soon after seedling emergence. Birch *et al.* (1998) confirmed that maize plants also acclimated to the temperature and that their phyllochron increased by 1.7°Cd per °C increase in the mean temperature within a 12–25°C mean temperature range with small inter-variety variability. The simultaneous acclimation of the phyllochron and the duration to PI was observed in three sorghum varieties sown in May and June 2001 in Bamako and Montpellier (Hemberger, 2001) (Supplementary Table 3). This adaptation to the colder mean temperature at seedling emergence in May and June in Montpellier did not happen in 2006 because of exceptionally high temperatures at seedling emergence. Thus, in Montpellier, the durations to PI were much longer in 2006 than in 2001 and in usual cropping seasons.

Acclimation of the duration to PI to the mean temperature has been previously observed. Thus, Yoshida (1981) reported a 1975 experiment in which four photoperiod-insensitive rice varieties with contrasting BVPs were grown in temperature-controlled glasshouses at 21, 24, 27, and 30°C. From 24°C upwards, the temperature had only a small effect on the duration to flowering, whereas this duration increased by 20 to 35 d when temperature decreased from 24 to 21°C. Yoshida (1981) hypothesised that below a threshold between 21 and 24°C, plants could no longer ensure the homeostatic control of the duration to flowering that they maintained between 24 and 30°C. Similarly, Roberts and Carpenter (1965) showed that the duration to flowering of one photoperiod-insensitive rice variety was also insensitive to the mean temperature between 30 and 35°C. In another seven rice varieties, the duration to PI increased by 10 to 20 d between 30°C and 35°C. In beans, Wallace *et al.* (1991) reported stable durations to the first flower between 23 and 26°C in 10 varieties grown at various altitudes in Guatemala and nearly stable durations between 18 and 24°C in a similar experiment with nine varieties in Colombia. These stable durations were accompanied by a nearly stable number of nodes below the first flower.

Thus, the continuous curvilinear response of the duration to flowering to the temperature often observed in growth chambers and used to support the photothermal model that currently predominates (Summerfield *et al.,* 1992), was not a unique relationship. Conversely, much data, including that presented in this paper, support the hypothesis of acclimation to temperature for the duration to flowering within a specific range of temperatures under natural lighting. Thus, plants actively create a discontinuous interval in the relationships between temperature and either the rate of leaf appearance or the duration to flowering. Precise information is lacking on the size and bounds of this range and on the level of homeostasis in the duration to flowering within this interval. However, in rice, the range would extend at least from 24 to 30°C (Roberts and Carpenter, 1965; Yoshida, 1985) and in sorghum at least from 19 to 31°C (Fig 6C); thus, they would cover the usual range of mean temperature for both crops. Assuming that acclimation can nearly stabilise the duration to flowering, as shown by Yoshida (1981) and in Fig. 5C, using days instead of thermal time for modelling the duration to PI in rice and sorghum appeared as a more accurate though perfectible choice.

### Crop-photoperiodismmodel2.0 perfectly ftied every variety patern for the duration to PI

In sorghum, no previous model could consider the longer duration of the vegetative phase observed for November and December sowings in many varieties grown in Bamako (Dingkuhn *et al.,* 2008) and other tropical locations (Bezot, 1963; Miller *et al.,* 1968). This was especially true for CSM 63-E, which was photoperiod-insensitive for May to October sowings and showed a reaction to the sowing date from November to April (Fig. 2B). The addition to the daylength effect in combined sensitivity to the daily change in sunrise time (dSR) and sunset time (dSS) fully resolved the point in every variety tested in Samanko without any temperature input. The combined effect of dSR and dSS successfully simulated the longer duration of the vegetative phase for sowings from July to October in the daylength-insensitive variety IRAT 204 (Fig. 2A).

In rice, the ORYZA model, now incorporated within the APSIM platform, simulates the duration of the PSP until PI (Bouman *et al.,* 2001). The RIDEV model accurately predicted the duration to flowering (not to flowering induction) of early to late varieties in northern Senegal where the model was calibrated and in Madagascar situated at a similar latitude after the adjunction of a temperature-acclimation coefficient (Dingkuhn *et al.,* 1995, 2008, 2015). To attain a good fit for the durations to PI recorded for rice in greenhouses, it was assumed that the glass of the greenhouses altered the sunlight effect on the duration to PI. This assumption agreed with the previous observation that the duration to PI was much shorter in plants grown in a greenhouse than in plants sown in the field in three sorghum varieties sown in December (Clerget *et al.,* 2012). This assumption was also supported by the monthly sowings of rice varieties in paddy fields in northern Senegal that showed long durations to flowering for sowings from October to February, especially for the variety Nipponbare present in both the African and Asian series (Supplementary Fig.4).

Thanks to knowledge attained by Borchert *et al.* (2005) and Goymann *et al.* (2012), the adjunction of the additive effects of the daily change in sunrise and sunset times to the existing photoperiodism model greatly improved its efficiency in predicting the sorghum and rice response to sowing date. An additional improvement of the model came from the hypothesis that plants could very differently combine the dSR and dSS cues either when dSS > dSR, before the summer solstice, or when dSS < dSR. Two sets of sensitivity parameters, SRs and SSs, were thus sought during the calibration. Such change in photoperiod-sensitivity at the summer solstice, when the daily change in the daylength switches from positive to negative, was already suspected (Curtis, 1968; Clerget *et al.,* 2004). However, evidence for the respective role of dSR and dSS needs to be determined.

For decades, the photoperiodic reaction has been divided as either quantitative or qualitative, depending on its strength, with no definition of the limit between the two classes (Thomas and Vince-Prue, 1997). Quantitative varieties were modelled with a simple linear relationship between the daylength and the duration to either flowering or PI (Major, 1980). Above a daylength threshold, PI was temporarily inhibited in models for qualitative varieties. The threshold could be either fixed (Carberry *et al.,* 1992) or moving (Folliard *et al,.* 2004; Dingkuhn *et al.* 2008). A unique linear relationship in the crop-photoperiodism model 2.0 regrouped all short-day plants into a continuum. The predicted duration to PI was sufficiently delayed by the very long duration of the daily-predicted duration of the PSP through high values of the sensitivity parameters (Fig. 7). Conversely, the concept of a memorised daily progress towards PI (Summerfield *et al.,* 1992) maintained its strength.

### Crop-photoperiodismmodel2.0 correclyt fited observations atother lauti des

In rice, once calibrated with data from lower latitudes, the observations conducted at a higher latitude were fitted well by the model (Fig. 3D&E). Conversely, in sorghum, the durations to PI predicted in Montpellier for May to July sowings were close to the exceptionally late durations observed in 2006 for Sariaso 10 and CSM 335 and too long for that of Souroukoukou (Fig. 3A-C). The model was calibrated with Bamako data, where monthly mean temperatures were less variable than those in Montpellier (Fig. 5B). In Montpellier, plants acclimated the duration to PI at exceptionally high temperatures during seedling emergence in 2006, whereas in most years, plants acclimated in response to the cool May temperatures. In regular years, June and July temperatures are much higher than the low temperature of acclimation; thus, plants would complete the short acclimated thermal time to PI in fewer days than that predicted by the model. The fortunate 2006 observations showed that in the same location at the same sowing date, the duration to PI was stable neither in days nor in thermal time because of the acclimation to the prevailing temperature at seedling emergence (Supplementary Table 3). It consequently appeared that the adaptation of the flowering time of tropical species, such as sorghum, to the temperate climate benefited from the usually cool temperature at seedling emergence.

Except for Siam 29, data are not yet sufficient to guarantee that the current variety parameters can be used at other latitudes than the latitude used for model calibration. It was disappointing to confirm that data on flowering time from greenhouse experiments could not be used for field crops, especially because monthly sowings over the entire year are not feasible at temperate latitudes. Such series were very helpful in the accurate parametrisation of Siam 29 and equatorial series would be very useful in sorghum. However, the results obtained for Siam 29 and from punctual comparisons in sorghum showed that the crop-photoperiodism model 2.0 could be parametrised to correctly predict the duration to PI at any latitude. Attaining the eight parameters of the crop-photoperiodism model 2.0 for each variety is a heavy task because the durations to PI of 12 consecutive monthly sowings in two tropical locations would likely be necessary.

## Supporting information

Computation of photoperiod

## ACKNOWLEDGEMENTS

Countless people participated in this long-term research, either through their support or work, and the authors are anonymously thankful to each of them. Many institutions participated in the funding and the implementation of the activities: CIRAD in France, ICRISAT in Mali, IRRI in the Philippines, CIAT and CORPOICA in Colombia, and INTA and the Federation of Cooperatives for Development (FECODESA) in Nicaragua.

