## Supplementary material for "Crop-photoperiodism model 2.0 for the panicle-initiation date of sorghum and rice that includes daily changes in sunrise and sunset times": Computation of photoperiod

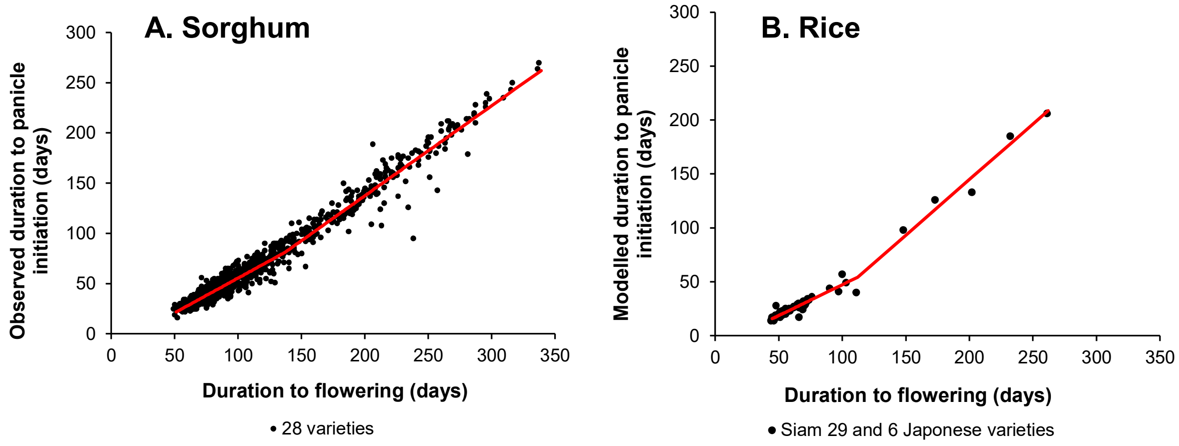


**Supplementary Fig. 1:** Relationships between the duration to panicle initiation and the duration to flowering in A) sorghum and B) rice.

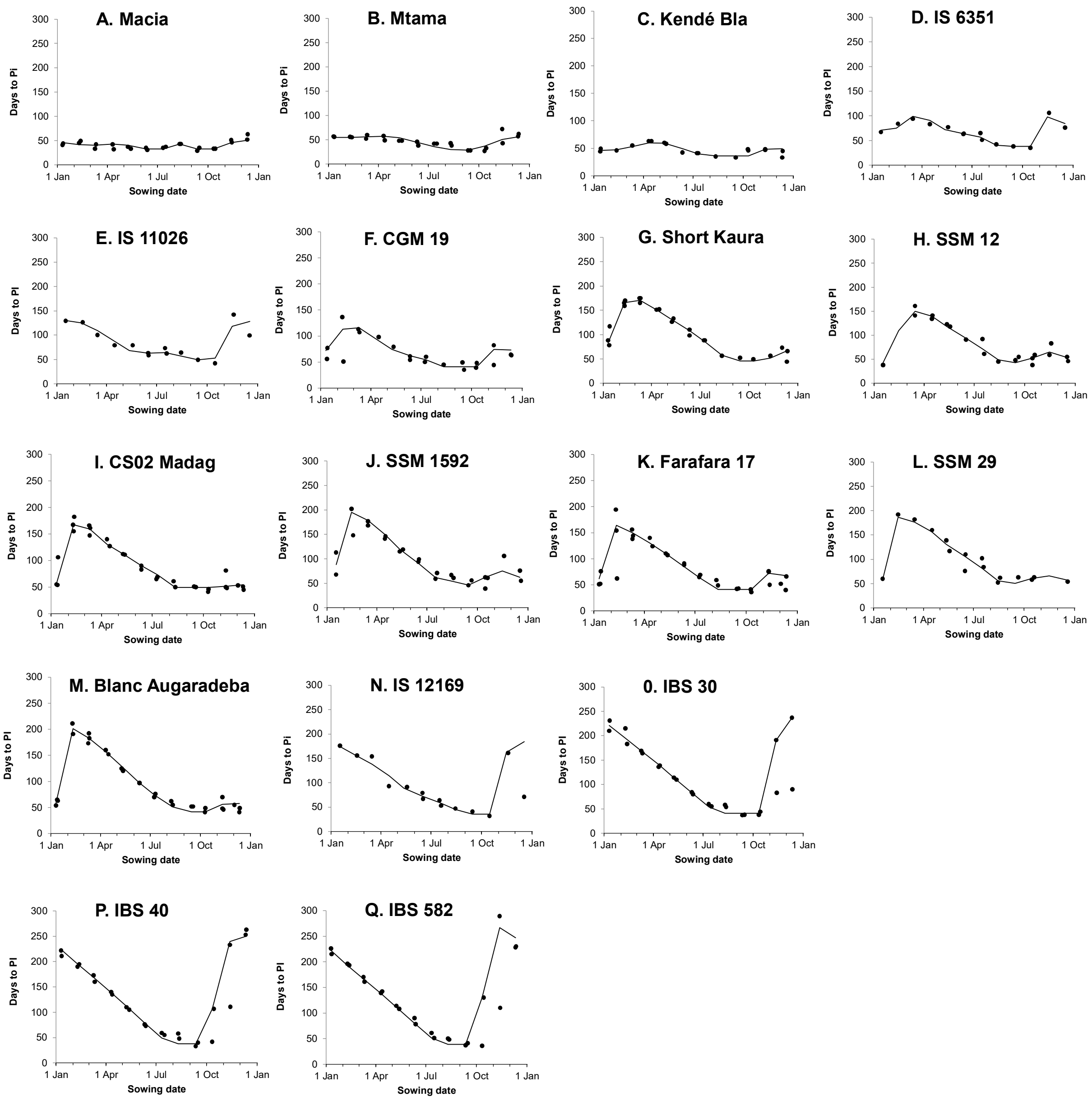


**Supplementary Fig. 2:** Two-year observations of the duration to panicle initiation by sowing month (●) and the values predicted by the model (─) calibrated on these observations in 17 sorghum varieties grown in Samanko (14°34'N).

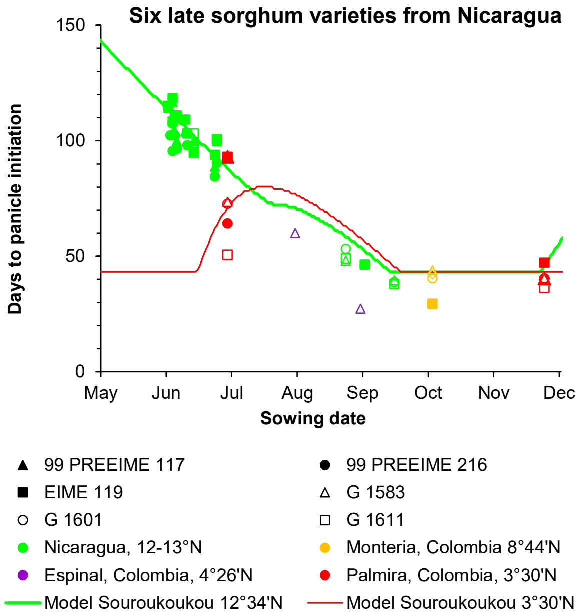


**Supplementary Fig. 3**: Duration to PI estimated from the observed duration to flowering of six Nicaraguan varieties sown from June to late-November at four contrasting latitudes, from 3°30'N to 13°N, compared with the predicted durations to PI of the variety Souroukoukou grown either in Samanko, Mali or in Palmira, Colombia.

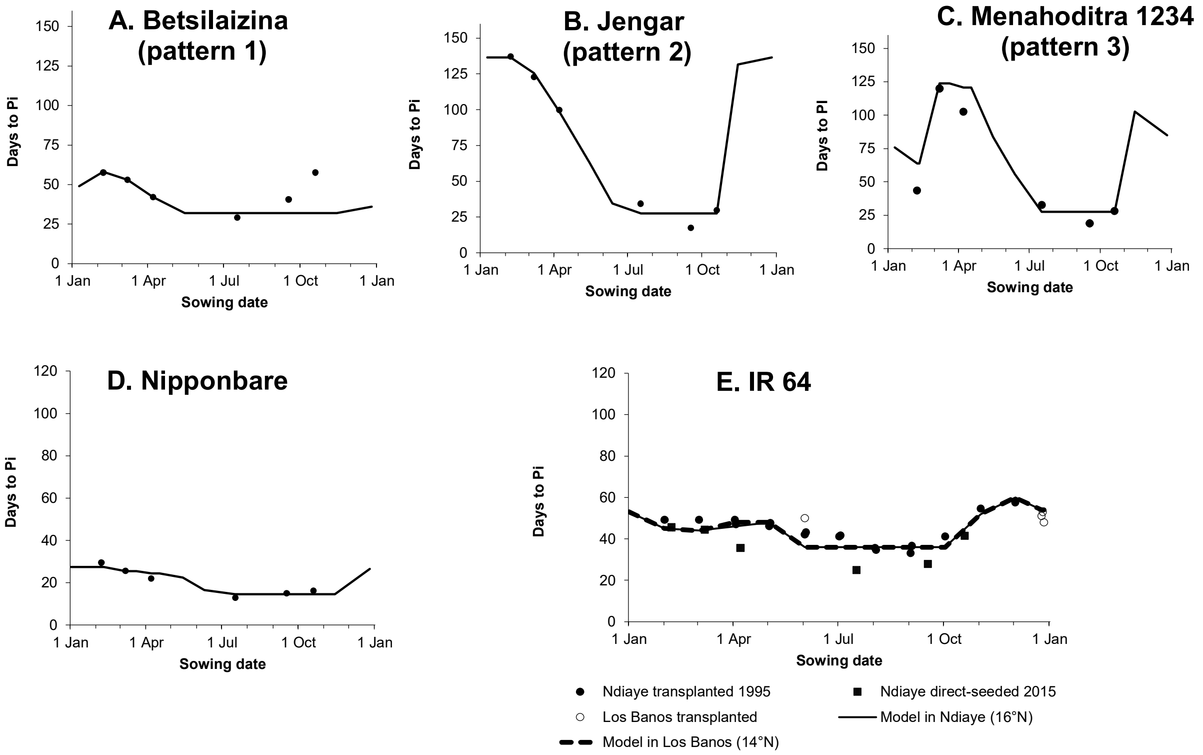


**Supplementary Fig. 4:** Observations of the duration to panicle initiation on contrasting sowing dates in Ndiaye (Senegal, 16°12'N) in 1991-1992 (■) and in 2009 (●) and in Los Baños (Philippines, 14°11'N) (○) of three rice varieties (A-C) representative for three different annual patterns (Dingkuhn et al. 2015), of Nipponbare (D) and the check variety IR 64 (E). Predictions from the crop-photoperiodism model 2.0 calibrated with Ndiaye data were plotted either for the latitude of Ndiaye (—) or for the latitude of Los Baños (---).

**Supplementary Table 1:** List of the 28 sorghum varieties monthly sown in Samanko, Mali.

| Genotype | Geographic origin | Race | Monthly sowing series |
|---|---|---|---|
| CSM335 | Mali | Guinea | All |
| IRAT 174 | Burkina Faso | Kafir-durra | 1 and 3 |
| Sariaso 10 | Burkina Faso | Caudatum | 1 and 3 |
| Blanc Augaradeba | Nigeria | Guinea | 2 |
| CGM 19 | Mali | Guinea-caudatum | 2 |
| CS 02 Mdg | Madagascar | Guinea margaritiferum | 2 |
| CSM 63E | Mali | Guinea | 2 |
| Farafara 17 | Nigeria | Guinea | 2 |
| IRAT 204 | Senegal | Caudatum | 2 |
| IS 15401 | Cameroon | Guinea-caudatum | 2 |
| Kaura D-12 | Nigeria | Durra-caudatum | 2 |
| Kendé Bla | Mali | Guinea margaritiferum | 2 |
| Ouéni | Mali | Guinea | 2 |
| Short Kaura | Nigeria | Durra-caudatum | 2 |
| Souroukoukou | Mali | Caudatum | 2 |
| IBS 19 | Tanzanie | Guinea | 3 |
| IBS 30 | Tanzanie | Guinea | 3 |
| IBS 40 | Tanzanie | Guinea | 3 |
| IBS 582 | Mozambique | Guinea | 3 |
| Macia | Zimbabwe | Caudatum | 3 |
| Mtama | Kenya | Caudatum | 3 |
| Sima | Zimbabwe | Guinea-caudatum | 3 |
| IS 6351 | India | Durra | 4 |
| IS 11026 | Ethiopia | Durra | 4 |
| IS 12169 | Ethiopia | Bicolor | 4 |
| SSM 12 | Cameroon | Durra | 4 |
| SSM 29 | Cameroon | Durra | 4 |
| SSM 1592 | Tchad | Durra | 4 |

**Supplementary Table 2:** List of the 16 sorghum varieties from the CIRAD Core Collection with duration to flowering over than 10 months when sown on 13 November 2007 in Samanko, Mali.

| Accession | Race | Origine | Flowering date | Duration to flowering (days) |
|---|---|---|---|---|
| IS 21519 | Guinea | Malawi | 12 Sept | 304 |
| IS 22893 | Guinea-Caudatum | Sudan | 4 Oct | 326 |
| IS 25499 | Caudatum | Burundi | 4 Oct | 326 |
| IS 10882 | Caudatum | Nigeria | 10 Oct | 332 |
| IS 24139 | Guinea | Tanzania | 10 Oct | 332 |
| SSM 1103 | Durra-Caudatum | Chad | 10 Oct | 332 |
| IS 23254 | Bicolor | Zimbabwe | 14 Oct | 336 |
| IS 10194 | Bicolor | Burkina Faso | 20 Oct | 342 |
| IS 16186 | Caudatum | Cameroun | 20 Oct | 342 |
| IS 10801 | Guinea-Caudatum | Chad | 24 Oct | 346 |
| IS 26554 | Guinea | Benin | 25 Oct | 347 |
| IS 2156 | Bicolor | Nigeria | 4 Nov | 357 |
| IS 17658 | Guinea | Ghana | 2 Nov | 355 |
| IS 23100 | Guinea | Tanzania | 3 Nov | 356 |
| IS 26457 | Guinea | Benin | 4 Nov | 357 |
| SSM 232 | Guinea | Senegal | 10 Nov | 363 |

**Supplementary Table 3:** Five developmental traits and ANOVA for three sorghum varieties simultaneously sown in Samanko, Mali (12°34'N) and in Montpellier, France (43°39'N) in May and June of 2001 and 2006.

| | | | Bamako | | | | | Montpellier | | | | |
|---|---|---|---|---|---|---|---|---|---|---|---|---|
| Variety | Year | Month | Phyllo1 (°Cd) | Phyllo2 (°Cd) | Days to PI | SumT to PI (°Cd) | Total Leaves | Phyllo1 (°Cd) | Phyllo2 (°Cd) | Days to PI | SumT to PI (°Cd) | Total Leaves |
| CSM 335 | 2001 | May | 44 | 80 | 81 | 1266 | 31.5 | 29 | 44 | | | 28.5 |
| | | June | 41 | 76 | 73 | 1053 | 29.3 | 32 | | | | |
| | 2006 | May | 40 | 88 | 83 | 1321 | 30.3 | 42 | 75 | 105 | 1235 | 31.5 |
| | | June | 44 | 72 | 71 | 1102 | 25.2 | 38 | 75 | 72 | 910 | 28 |
| IRAT 174 | 2001 | May | 44 | 52 | 79 | 1236 | 33.8 | 29 | 44 | | | |
| | | June | 41 | 60 | 71 | 1021 | 31.2 | 31 | | | | |
| | 2006 | May | 40 | 62 | 75 | 1209 | 33 | 40 | 51 | 37.4 | 1246 | 37.4 |
| | | June | 49 | 62 | 57 | 886 | 27.4 | 39 | 53 | 30 | 955 | 30 |
| Sariaso 10 | 2001 | May | 47 | | 54 | 904 | 28 | 35 | - | 50 | 502 | 20.6 |
| | | June | 41 | 65 | 50 | 735 | 27.6 | 33 | - | 43 | 492 | 20.1 |
| | 2006 | May | 37 | 64 | 60 | 978 | 26.4 | 41 | 53 | 82 | 984 | 29.6 |
| | | June | 50 | 66 | 57 | 888 | 25.6 | 37 | 52 | 54 | 731 | 25.3 |

| **ANOVA** | | Phyllo1 | Phyllo2 | Days to PI | SumT to PI | Total Leaves |
|---|---|---|---|---|---|---|
| Factor | DF | | | F | | |
| Location | 1 | 22.51 *** | 0.88 ns | 0.85 ns | 9.22 ** | 11.38 ** |
| Year(location) | 2 | 3.94 * | 0.05 ns | 1.25 ns | 4.00 * | 4.56 * |
| Month(year) | 1 | 90.04 *** | 5.23 * | 1.73 ns | 13.77 ** | 3.02 ns |
| Month*location(year) | 2 | 15.43 *** | 0.13 ns | 0.14 ns | 0.35 ns | 0.49 ns |

**Supplementary Table 4:** Parameters of the panicle initiation model and data fitting of 17 sorghum varieties monthly sown in Bamako. When data from several locations were available, they were used for either calibrating (C in Parm) or validating (Val) the model. Data fitting was computed either for all data from twelve sowing months or for data from only eight sowing months, from March to October.

| | | | Parameters | | | | | | | | Data fitting | | | |
|---|---|---|---|---|---|---|---|---|---|---|---|---|---|---|
| | | | BVP | $P_{b1}$ | Ps | $SR_{s1}$ | $SS_{s1}$ | $P_{b2}$ | $SR_{s2}$ | $SS_{s2}$ | 12 months | | March-October | |
| Variety | Location | Parm | days | h | day $h^{-1}$ | $day^2$ $s^{-1}$ | $day^2$ $s^{-1}$ | h | $day^2$ $s^{-1}$ | $day^2$ $s^{-1}$ | $R^2$ | RMSD | $R^2$ | RMSD |
| Macia | Bamako | C | 31.81 | - | - | -0.1867 | 0.6860 | 12.04 | -5.392 | -0.149 | 0.50 | 5.6 | 0.02 | 5.7 |
| Mtama | Bamako | C | 27 | 8.24 | 4.94 | -0.156 | 0.4561 | - | - | - | 0.64 | 7.1 | 0.65 | 6.7 |
| IS 11026 | Bamako | C | 48.0 | 10.46 | 19.14 | -1.1769 | 1.7042 | 11.76 | -1.697 | 0.4537 | 0.85 | 11.8 | 0.81 | 7.4 |
| CGM 19 | Bamako | C | 40.0 | 11.88 | 91.63 | -0.4585 | 1.2502 | 12.02 | -0.305 | 0.6749 | 0.62 | 15.2 | 0.86 | 10.3 |
| Kendé Bla | Bamako | C | 35.0 | 11.22 | 12.25 | 0.0094 | 0.3455 | 11.35 | 0.1797 | 1.5112 | 0.84 | 3.5 | 0.86 | 3.9 |
| IS 12169 | Bamako | C | 34.23 | 11.05 | 60.19 | -1.8901 | 3.5791 | 11.41 | -2.877 | 1.3361 | 0.67 | 31.3 | 0.92 | 9.4 |
| IS 6351 | Bamako | C | 37.10 | 11.35 | 29.13 | -0.0782 | 1.9977 | 12.13 | -0.512 | 0.2501 | 0.92 | 5.7 | 0.95 | 4.8 |
| CS02 Madagascar | Bamako | C | 48.39 | 12.0 | 194.17 | -0.1423 | 0.1881 | 12.44 | 2.0538 | 2.5588 | 0.93 | 12.6 | 0.98 | 6.3 |
| Farafara 17 | Bamako | C | 40.0 | 11.90 | 174.20 | -0.2483 | 1.0825 | 12.54 | 0.7235 | 4.2292 | 0.75 | 22.2 | 0.98 | 7.1 |
| Short Kaura | Bamako | C | 44.77 | 11.78 | 146.23 | -0.0237 | 0.9415 | 9.65 | -0.706 | 0.2716 | 0.96 | 9.4 | 0.99 | 4.6 |
| SSM 12 | Bamako | C | 41.73 | 11.86 | 129.21 | 0.2485 | 0.4702 | - | - | - | 0.95 | 9.1 | 0.95 | 9.5 |
| SSM 1592 | Bamako | C | 44.45 | 11.77 | 174.88 | 0.1501 | 0.8106 | 12.11 | 6.8561 | 1.7774 | 0.92 | 14.3 | 0.97 | 8.8 |
| SSM 29 | Bamako | C | 49.59 | 11.87 | 199.84 | 0.1135 | 0.3995 | - | - | - | 0.94 | 10.8 | 0.92 | 11.9 |
| Blanc Augaradeba | Bamako | C | 41.54 | 11.87 | 227.10 | -0.2584 | 0.4430 | 12.18 | 6.22 | 1.4925 | 0.98 | 7.5 | 0.99 | 6.1 |
| IBS 30 | Bamako | C | 40 | 11.39 | 161.21 | -2.360 | 3.6242 | - | - | - | 0.74 | 38.0 | 0.99 | 6.2 |
| IBS 40 | Bamako | C | 36.79 | 11.19 | 152.76 | -1.3485 | 4.8213 | - | - | - | 0.82 | 33.4 | 0.96 | 7.5 |
| IBS 582 | Bamako | C | 37.45 | 10.48 | 101.13 | -3.3807 | 4.5315 | - | - | - | 0.83 | 26.7 | 0.77 | 22.8 |

**Supplementary Table 5:** Parameters of the panicle initiation model and data fitting of A) three rice varieties grown in Ndiaye and representative of the three observed patterns of photoperiod-sensitivity (Dingkuhn 2015), and of B) one rice varieties grown in Ndiaye and in Los Baños. When data from several locations were available, they were used for either calibrating (C in Parm) or validating (Val) the model. Data fitting was computed either for all data from twelve sowing months or for data from only eight sowing months, from March to October.

| | | | | Parameters | | | | | | | | Data fitting | | | |
|---|---|---|---|---|---|---|---|---|---|---|---|---|---|---|---|
| | | | | BVP | $P_{b1}$ | Ps | $SR_{s1}$ | $SS_{s1}$ | $P_{b2}$ | $SR_{s2}$ | $SS_{s2}$ | 12 months | | March-October | |
| Variety | Model type | Location | Parm | days | h | day $h^{-1}$ | $day^2\ s^{-1}$ | $day^2\ s^{-1}$ | h | $day^2\ s^{-1}$ | $day^2\ s^{-1}$ | $R^2$ | RMSD | $R^2$ | RMSD |
| **A. One location** | | | | | | | | | | | | | | | |
| Betsilaizina | Field | Ndiaye | C | 29.88 | - | - | -0.463 | 0.6502 | 12.45 | 0.7706 | -0.463 | 0.92 | 3.1 | 0.90 | 3.6 |
| Jengar | Field | Ndiaye | C | 22.25 | 12.31 | 78.86 | -1.582 | 3.921 | 10.80 | -0.618 | 0.1606 | 0.99 | 3.6 | 0.98 | 4.4 |
| Menahoditra 1234 | Field | Ndiaye | C | 21.29 | 12.12 | 186.79 | -0.361 | 0.5656 | 10.69 | -6.66 | 0.2953 | 1.00 | 2.8 | 0.99 | 3.4 |
| **B. Two locations** | | | | | | | | | | | | | | | |
| IR 64 | Field | Ndiaye 1995 | C | 34 | - | - | -0.089 | 0.7836 | 11.39 | 0.673 | 0.1343 | 0.87 | 2.5 | 0.90 | 2.0 |
| | ‘’ | Ndiaye 2015 | Val | | | | | | | | | 0.15 | 9.5 | - | - |
| | ‘’ | Los Baños | Val | | | | | | | | | - | 3.9 | | |
